# Hyperactivation of β-catenin rewires mitochondrial fuel selection in liver cancer cells

**DOI:** 10.64898/2026.10.18.759986

**Authors:** Luis Cedeño-Rosario, Queren A. Alcantara, Nicole A. Sierra, Praveena Prasad, Jillian Wright, Geanette Lam, Nimo Abdi, Jaide Florence, Alex J. Bott, Katarina Heyden, Joshua C. Youngs, Álvaro J. Narbona-Pérez, Lorenzo Gastelum Mena, Monika A. Makurath, Jesse N. Velasco-Silva, Mi-Young Jeong, Sarah Fogarty, Gregory S. Ducker, Jennifer Lippincott-Schwartz, Melanie R. McReynolds, Jared Rutter

## Abstract

Metabolic state is tightly linked to cell fate, and expression of the mitochondrial pyruvate carrier (MPC), which imports pyruvate into mitochondria, is a robust marker of it. The transcriptional regulators that set metabolic state, however, remain poorly defined. In the liver, β-catenin is one of the determinants of metabolic zonation, and activating *CTNNB1* mutations are frequent in hepatocellular carcinoma. Here we show that β-catenin regulates metabolic state in liver cancer cells. An unbiased CRISPR screen that uses endogenous *MPC1* transcript abundance as the marker identifies *CTNNB1*. Activating β-catenin pharmacologically or genetically reduces MPC1 and MPC2 transcript and protein levels, and MPC expression is also low in β-catenin-driven liver tumors in vivo. Beyond the MPC, β-catenin activation impairs assembly of respiratory Complexes III and IV, reduces mitochondrial DNA content and respiratory capacity, elongates mitochondrial networks and remodels the lipidome. Stable isotope tracing shows reduced entry of glucose- and lactate-derived carbon into the TCA cycle alongside increased oxidation of palmitate and oleate. These findings identify β-catenin as a regulator of metabolic state that drives a fuel switch from carbohydrate to fatty acid oxidation in liver cancer cells.

## INTRODUCTION

As cells transition through development, respond to environmental cues or pathologically transform, they frequently change metabolic state. Over the past 15 years, we have learned through causation-directed experiments that this metabolic state has profound impacts on cell fate determination^1–9^. Metabolic state is a complex concept that includes mechanisms of energy extraction and generation of ATP, production of the necessary intermediates for macromolecule synthesis, and many other processes^10^. We and others have shown that one particular metabolic node plays a particularly important role in determining the fate and function of cells, and that is the pyruvate-lactate metabolic axis^11–13^.

Typically, cells convert much of their imported glucose into pyruvate in the cytosol through the reactions of glycolysis. This pyruvate can then be converted to lactate and exported, which is the classic metabolic program of cancer cells and other rapidly proliferating cell types. Alternatively, pyruvate can be transported into mitochondria to feed the tricarboxylic acid (TCA) cycle. The mitochondrial pyruvate carrier (MPC), a heterodimer composed of the MPC1 and MPC2 proteins, is the transporter that allows pyruvate to enter the mitochondrial matrix^14–17^. Because it sits at the junction between glycolysis and mitochondrial oxidation, the MPC occupies a point of metabolic bifurcation, and expression of the MPC is a robust marker of cellular metabolic state. Expression of the MPC genes is low in intestinal stem cells and in colon cancer cells at both the mRNA and protein levels, and this low expression promotes proliferation and oncogenesis^5,7^. More generally, low MPC expression in cultured cells correlates with a stem cell-like gene expression signature^5,6^. These data convinced us that the expression of the MPC genes, particularly *MPC1*, is a faithful marker of metabolic state and we were motivated to use this marker as a handle to understand the transcriptional drivers of metabolic state.

Our previous work showed that MPC expression is also negatively correlated with the expression of canonical targets of the Wnt/β-catenin signaling pathway, and that MPC inactivation promotes tumor initiation in the intestine^7^. The Wnt/β-catenin pathway maintains the balance between stemness and differentiation^18–20^. In the canonical pathway, Wnt ligand binding represses the β-catenin destruction complex, composed of APC, GSK3, CK1, PP2A and β-TrCP, allowing β-catenin to accumulate and translocate to the nucleus^21^. Nuclear β-catenin then regulates the expression of downstream target genes, including MYC and cyclin D1, through the TCF/LEF transcription factors^22^.

In the liver, Wnt/β-catenin signaling has an additional role as one of the principal determinants of metabolic zonation. β-catenin activity is restricted to a narrow perivenous zone, where glycolytic and lipogenic programs predominate, while oxidative and gluconeogenic programs are confined to the periportal zone^23,24^, although recent work indicates that hepatocyte metabolic states are more finely structured than this classical two-zone model implies^25^. Activating mutations in *CTNNB1*, the gene encoding β-catenin, are among the most frequent oncogenic events in hepatocellular carcinoma (HCC)^26^, and β-catenin-activated HCC tumors have been shown to be atypically less glycolytic and more dependent on fatty acid oxidation^27^. Whether and how β-catenin drives the switch in fuel preference in these tumors has remained unclear.

We set out to define the transcriptional drivers of metabolic state using *MPC1* as a marker gene. Genetic screens led us to the observation that β-catenin is a negative regulator of the expression of the MPC genes in liver cancer cells. We further show that activation of β-catenin, either pharmacologically or genetically, decreases MPC1 and MPC2 at both the mRNA and protein levels. Low MPC expression is also a feature of β-catenin-driven liver tumors in vivo. In addition, activation of β-catenin alters the metabolic program more broadly, including impacts on mitochondrial respiratory capacity, mitochondrial morphology, and the lipidome. Using stable isotope tracing, we find that β-catenin activation reduces the contribution of glucose- and lactate-derived carbon to the TCA cycle while increasing the oxidation of long-chain fatty acids. Together, these results demonstrate that β-catenin signaling rewires the metabolic program of liver cancer cells by engineering a fuel switch from carbohydrate to fat oxidation.

## RESULTS

### β-catenin is a repressor of MPC expression

To identify transcriptional regulators of metabolic state, we conducted a fluorescence-activated cell sorting (FACS)-based genetic screening assay (TRoUT-FISH) to discover genes that positively or negatively regulate *MPC1* abundance^28^. This approach measures the abundance of an endogenous transcript directly by RNA FISH, and therefore does not require an exogenous reporter or modification of the target locus^28^. We infected JHH7 liver cancer cells with a CRISPR sgRNA library designed against genes involved in transcriptional and epigenetic regulation, performed RNA FISH against *MPC1*, and sorted the cells into *MPC1*-high and *MPC1*-low populations, after which sgRNA abundance in each population was determined by sequencing (Fig. 1a). Interestingly, we found that *CTNNB1* (the gene that encodes the β-catenin protein) negatively regulated *MPC1* expression, such that its deletion caused an increase in *MPC1* mRNA abundance (Fig. 1b). Moreover, we also identified *TCF7* and *LEF1* (additional components of the Wnt/β-catenin signaling pathway) whose deletion had the same effect (Fig. 1b), consistent with prior CRISPR screens that recovered coherent sets of Wnt pathway components in cancer cells^29^. Based on these results, we hypothesized that activation of the Wnt/β-catenin pathway represses MPC gene expression.

**Fig. 1.**
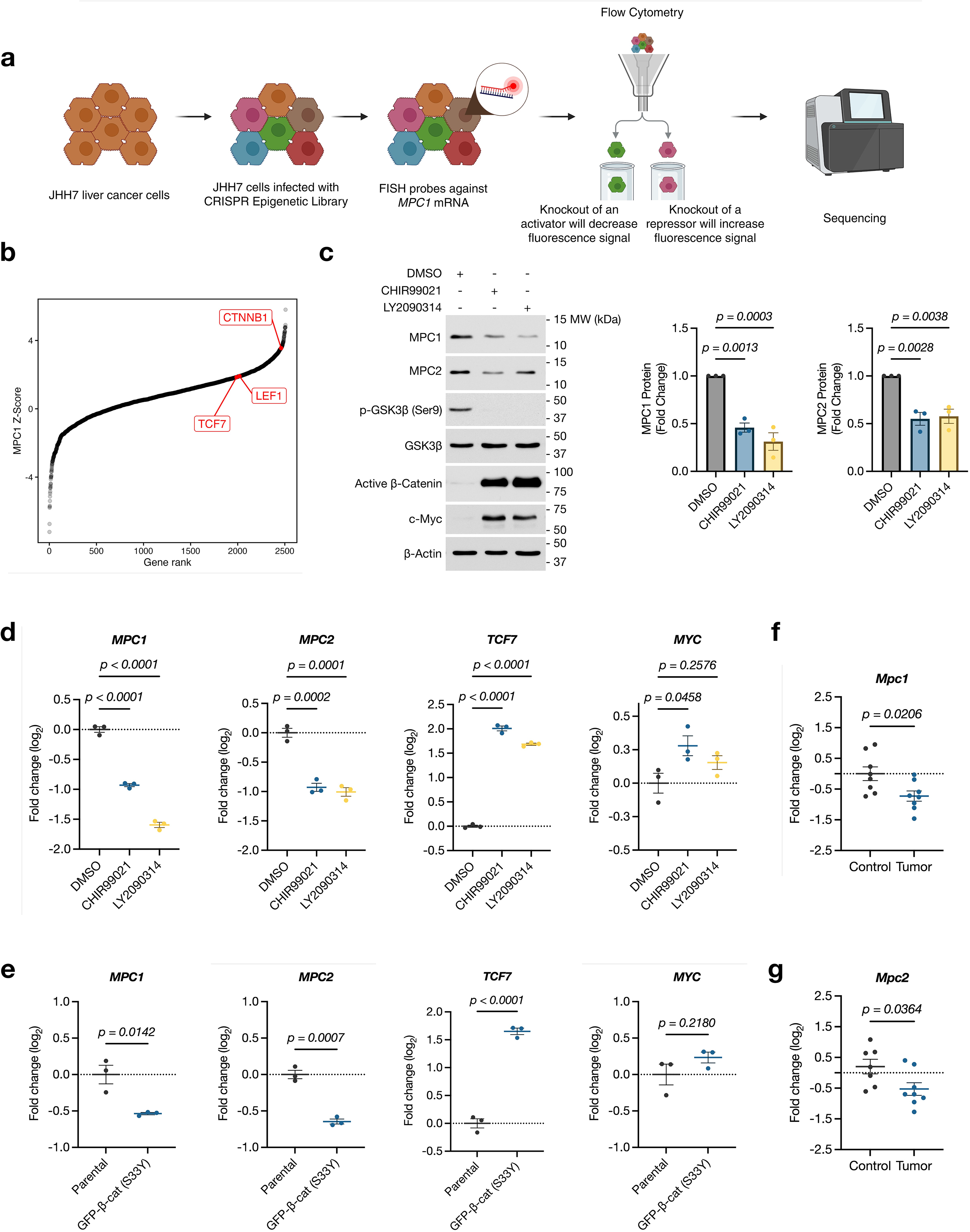
β-catenin represses MPC expression in liver cancer cells. **a**, Schematic of the TRoUT-FISH screen. JHH7 liver cancer cells were transduced with a pooled CRISPR epigenetic library (Human Epigenetic Knockout Library) and selected, fixed, and subjected to RNA FISH against *MPC1*. Cells were sorted by flow cytometry into MPC1-high and MPC1-low populations and sgRNA abundance was determined by sequencing. **b**, Gene-level results of the screen, ranked by z-score derived from MAUDE comparing the MPC1-high and MPC1-low populations; positive values indicate enrichment in the MPC1-high population, that is, genes whose deletion increases *MPC1* abundance. *CTNNB1, TCF7* and *LEF1* are highlighted. **c**, Immunoblot of whole-cell extracts from JHH7 cells treated with DMSO, 20 µM CHIR99021 or 5 µM LY2090314 for 48 h, probed for MPC1, MPC2, p-GSK3β (Ser9), total GSK3β, active β-catenin, c-Myc and β-actin. A representative blot is shown. Right, densitometric quantification of MPC1 and MPC2 normalized to β-actin and expressed as fold change relative to DMSO. Data are mean ± s.e.m. of *n* = 3 independent biological replicates; significance was assessed by One-way ANOVA with Dunnett correction. **d**, RT–qPCR of *MPC1*, *MPC2*, *TCF7* and *MYC* in JHH7 cells treated as in **c**. Data are log₂ fold change relative to DMSO, mean ± s.e.m. of *n* = 3 independent biological replicates; One-way ANOVA with Dunnett correction. **e**, RT–qPCR of the indicated transcripts in parental JHH7 cells and JHH7 cells stably expressing GFP-tagged β-catenin (S33Y). Data are log₂ fold change relative to parental cells, mean ± s.e.m. of *n* = 3 independent biological replicates; two-tailed unpaired Student’s *t*-test. **f**,**g**, RT–qPCR of *Mpc1* (**f**) and *Mpc2* (**g**) in liver tumors from mice carrying Sleeping Beauty transposons encoding human *CTNNB1* and *MYC*, compared with control (healthy liver tissue). Data are log₂ fold change relative to control, mean ± s.e.m.; *n* = 8 control and 8 tumor samples; two-tailed unpaired Student’s *t*-test. Exact *p* values are shown in each panel.

To test whether activation of β-catenin reduces MPC1 and MPC2 protein abundance, we activated β-catenin by treating JHH7 cells with two structurally distinct GSK3 inhibitors (CHIR99021 or LY2090314). After treatment, we analyzed whole cell extracts by SDS-PAGE and immunoblotting. The activation of β-catenin was confirmed by the profound increase in β-catenin that is un-phosphorylated at Ser45, a readout of stabilized and functionally active protein. c-Myc protein was also increased, consistent with the loss of GSK3-dependent phosphorylation of c-Myc at Thr58, which primes it for ubiquitin-mediated degradation^30,31^ (Fig. 1c). We found that treatment with either GSK3 inhibitor decreased MPC1 and MPC2 protein when compared to vehicle (Fig. 1c). These results suggest that activation of β-catenin is sufficient to decrease MPC1 and MPC2 protein abundance.

To investigate whether activation of β-catenin also reduced MPC1 and MPC2 expression at the mRNA level, we treated JHH7 cells with the GSK3 inhibitors mentioned above and analyzed transcript abundance via RT-qPCR. The activation of β-catenin was confirmed by the increase in *TCF7*, a direct downstream target of β-catenin^32^ (Fig. 1d). We found that *MPC1* and *MPC2* mRNA levels were decreased when compared to control (Fig. 1d), including in two additional liver cancer cell lines, Huh-7 and Hep3B (Extended Data Fig. 1a,b). In contrast, *c-MYC* transcript was elevated in Huh-7, modestly elevated in JHH7 cells and not elevated in Hep3B cells (Fig. 1d and Extended Data Fig. 1a,b), indicating that *c-MYC* mRNA induction is not a universal readout of pathway activation in this setting.

Because GSK3 has substrates beyond β-catenin^30^, we next asked whether stabilizing β-catenin alone is sufficient to repress *MPC1* and *MPC2* gene expression. We generated JHH7 cells stably expressing GFP-tagged β-catenin carrying the activating S33Y substitution, which escapes degradation by the destruction complex. These cells showed induction of *TCF7* consistent with hyperactive β-catenin. They also showed reduced *MPC1* and *MPC2* mRNA abundance (Fig. 1e).

To determine whether MPC repression also accompanies β-catenin-driven tumorigenesis in vivo, we collected liver tissue from a Sleeping Beauty transposon mouse model of HCC in which human *CTNNB1* and *MYC* are co-expressed in hepatocytes (Extended Data Fig. 1c). We found that both *Mpc1* and *Mpc2* transcripts were reduced in tumor tissue when compared to control liver (Fig. 1f,g). Because tumors in this model are driven by both β-catenin and MYC, these data do not isolate the contribution of β-catenin, but they establish that low MPC expression is a feature of β-catenin-driven liver tumors in vivo.

Taken together, these results identify β-catenin as a negative regulator of MPC1 and MPC2, demonstrate that activating β-catenin pharmacologically or genetically is sufficient to decrease MPC expression in liver cancer cells, and show that low MPC expression is a feature of β-catenin-driven HCC in vivo.

### β-catenin activation reduces mitochondrial respiratory capacity

Inspired by this new understanding that β-catenin represses expression of the MPC genes and potentially pyruvate oxidation, we sought to more comprehensively define how β-catenin activation affects mitochondrial function more broadly. Therefore, we performed RNA sequencing on parental JHH7 cells and JHH7 cells expressing β-catenin (S33Y). Gene set enrichment analysis showed positive enrichment of oxidative phosphorylation among the most significantly enriched hallmark gene sets in β-catenin (S33Y) cells (Fig. 2a), and analysis against MitoCarta gene sets showed enrichment of OXPHOS, Complex I and Complex V categories (Extended Data Fig. 2a). Consistent with this, nuclear-encoded subunits of all five respiratory complexes were nearly universally upregulated (Fig. 2b).

**Fig. 2.**
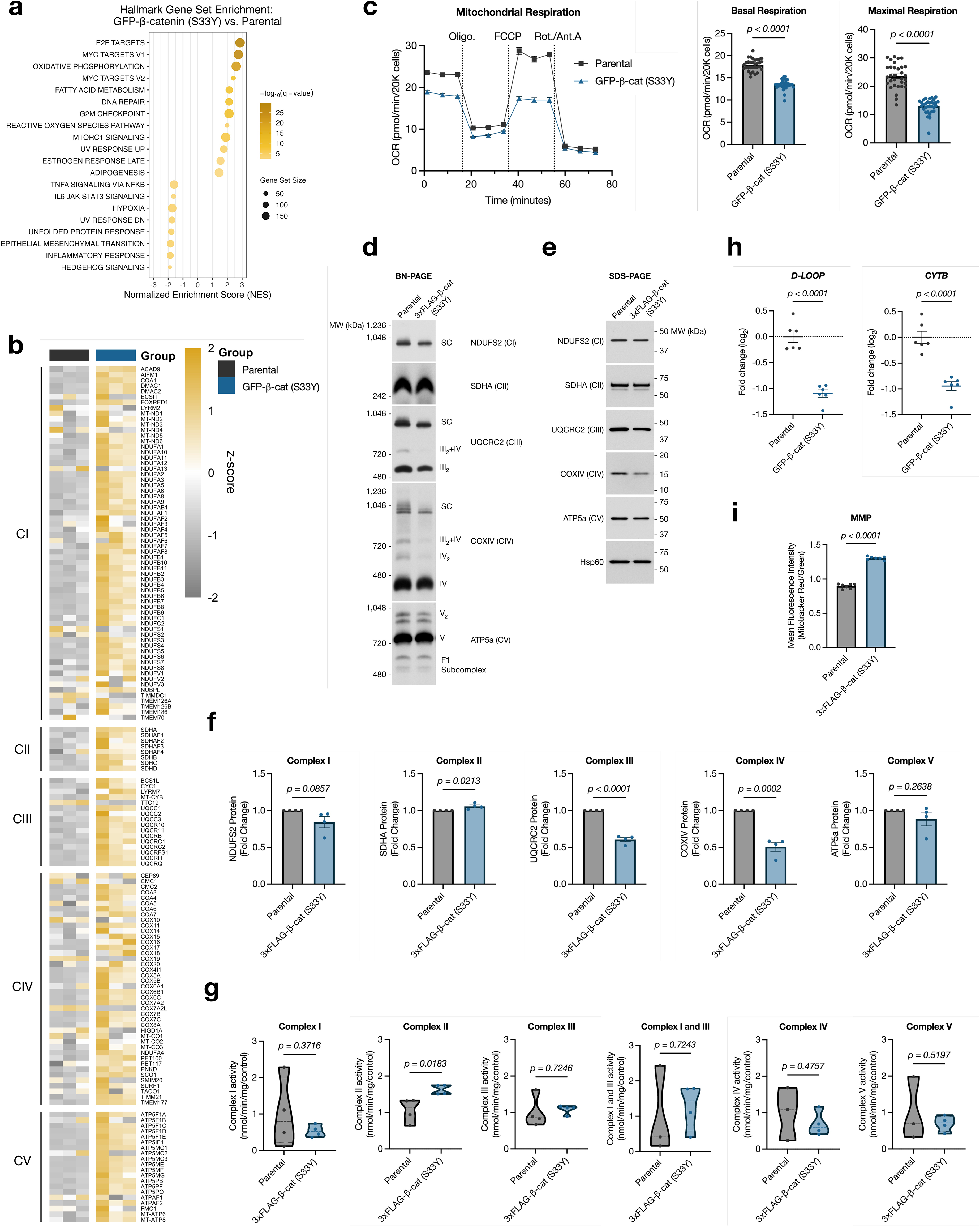
β-catenin activation compromises respiratory complex assembly and mitochondrial respiration. **a**, Hallmark gene set enrichment analysis of RNA sequencing data from JHH7 cells expressing β-catenin (S33Y) versus parental JHH7 cells (*n* = 3 independent biological replicates per group). Gene sets are ranked by normalized enrichment score (NES); dot size indicates gene set size and color indicates −log₁₀(*q* value). **b**, Heatmap of genes encoding subunits of respiratory Complexes I–V in parental and β-catenin (S33Y) cells, shown as *z*-scores of DESeq2-normalized counts, with genes ordered by respiratory complex; *n* = 3 independent biological replicates per group. **c**, Oxygen consumption rate (OCR) measured by Seahorse extracellular flux analysis in parental and β-catenin (S33Y) cells. Left, representative trace showing sequential injection of oligomycin, FCCP and rotenone/antimycin A. Right, basal and maximal respiration. Data are mean ± s.e.m. of *n* = 4 independent experiments, each comprising 8 technical replicate wells; statistical significance was assessed on the experimental means by two-tailed unpaired Student’s *t*-test. OCR was normalized to 20,000 cells. **d**, Blue native PAGE followed by immunoblotting for NDUFS2 (Complex I), SDHA (Complex II), UQCRC2 (Complex III), COXIV (Complex IV) and ATP5a (Complex V) in parental and β-catenin (S33Y) cells. Supercomplexes (SC), individual complexes and subassemblies are indicated. A representative blot of *n* = 4 independent experiments is shown. **e**, SDS-PAGE followed by immunoblotting for the same subunits, with Hsp60 as a loading control. A representative blot is shown. **f**, Densitometric quantification of the immunoblots in **e**, normalized to Hsp60 and expressed as fold change relative to parental cells. Data are mean ± s.e.m. of *n* = 4 independent biological replicates; two-tailed unpaired Student’s *t*-test. **g**, Enzymatic activity of Complexes I, II, III, I and III, IV and V in parental and β-catenin (S33Y) cells, expressed as nmol min⁻¹ mg⁻¹ relative to control. Violin plots show the distribution of *n* = 4 independent biological replicates; two-tailed unpaired Student’s *t*-test. **h**, Mitochondrial DNA copy number in parental and β-catenin (S33Y) cells, assessed by qPCR of the *D-LOOP* and *CYTB* loci normalized to nuclear DNA. Data are log₂ fold change relative to parental cells, mean ± s.e.m. of *n* = 6 independent biological replicates; two-tailed unpaired Student’s *t*-test. **i**, Mitochondrial membrane potential (MMP) in parental and β-catenin (S33Y) cells, measured by flow cytometry as the ratio of MitoTracker Red to MitoTracker Green mean fluorescence intensity. Data are mean ± s.e.m. of *n* = 7 independent biological replicates; two-tailed unpaired Student’s *t*-test. Exact *p* values are shown in each panel.

To determine whether this transcriptional program is reflected in mitochondrial function, we measured oxygen consumption rates using a Seahorse extracellular flux analyzer in cells cultured in high-glucose medium. In contrast to the mRNA abundance data, β-catenin (S33Y) cells exhibited decreased basal and maximal respiration when compared to parental cells (Fig. 2c), as well as decreased ATP production-coupled respiration, proton leak and spare respiratory capacity (Extended Data Fig. 2b). Thus, activation of β-catenin reduces mitochondrial respiration despite an increase in the expression of the genes encoding the respiratory machinery.

To understand the basis of this discrepancy, we examined the assembly and abundance of the respiratory complexes by blue native PAGE and SDS-PAGE, respectively, followed by immunoblotting against OXPHOS subunits. β-catenin (S33Y) cells showed decreased Complex III and Complex IV, as well as decreased abundance of the supercomplex form of Complex I, consistent with its physical association with Complexes III and IV. We did not observe assembly defects in Complex II or Complex V (Fig. 2d), consistent with the general consensus that they do not participate in respiratory supercomplexes^33^. In parallel, UQCRC2 (Complex III) and COXIV (Complex IV) protein levels were decreased, whereas SDHA (Complex II) was slightly increased and NDUFS2 (Complex I) and ATP5a (Complex V) were unchanged (Fig. 2e,f), suggesting that Complexes III and IV are both less abundant and less efficiently assembled in β-catenin (S33Y) cells. We next measured the enzymatic activity of each complex and, with the exception of a modest increase in Complex II activity, found that they were not significantly different between parental and β-catenin (S33Y) cells (Fig. 2g). Together, these data suggest that the decrease in mitochondrial respiration upon β-catenin activation might be due to defects in the assembly and abundance of Complexes III and IV, and not to a loss of catalytic activity of the individual complexes.

We therefore asked whether mitochondrial genome content was also affected. Quantification of mitochondrial DNA relative to nuclear DNA revealed that both *D-LOOP* and *CYTB* were reduced in β-catenin (S33Y) cells (Fig. 2h), indicating a decrease in mtDNA copy number. Paradoxically, we found that mitochondrial membrane potential was increased in β-catenin (S33Y) cells when compared to parental cells (Fig. 2i and Extended Data Fig. 2c).

Taken together, these results demonstrate that activation of β-catenin decreases mitochondrial respiratory capacity in liver cancer cells. This reduction occurs despite increased expression of nuclear-encoded respiratory chain genes and largely preserved activities of the individual complexes. It is instead associated with disrupted assembly and reduced abundance of Complexes III and IV, together with reduced mtDNA copy number. These combined data raise the possibility that the transcriptional upregulation of the respiratory machinery represents a compensatory response to impaired assembly and diminished mitochondrial genome content.

### β-catenin promotes mitochondrial network interconnectivity and reduces lipid droplet content

Changes in mitochondrial respiratory capacity are frequently accompanied by remodeling of mitochondrial network architecture, and mitochondrial elongation has been described as an adaptive response that sustains organelle function under metabolic stress^34,35^. We therefore examined mitochondrial morphology by immunofluorescence against TOMM20 in parental and β-catenin (S33Y) cells. β-catenin (S33Y) cells showed elongated, highly interconnected mitochondrial networks when compared to the more fragmented networks observed in parental cells (Fig. 3a). Quantitative morphometric analysis confirmed these observations: mitochondria in β-catenin (S33Y) cells exhibited increased area, perimeter, aspect ratio and form factor, together with increased branches, branch junctions, branch end points, mean branch length and total branch length per mitochondrion (Fig. 3b). Mean branch diameter was unchanged (Fig. 3b), indicating that these changes reflect network elongation and interconnectivity rather than an increase in mitochondrial width.

**Fig. 3.**
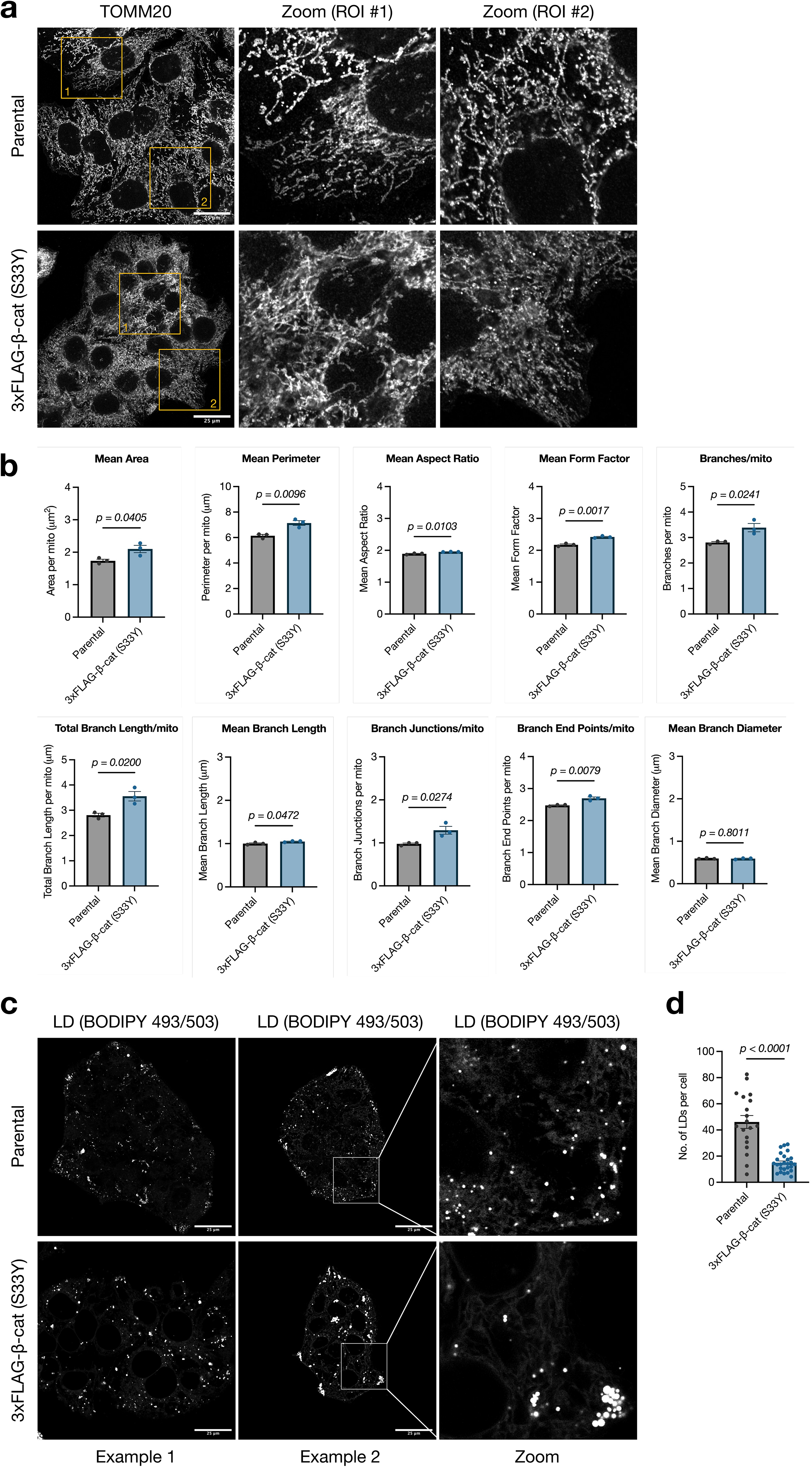
β-catenin activation drives mitochondrial network elongation and reduces lipid droplet number. **a**, Representative immunofluorescence images of mitochondrial networks in parental and β-catenin (S33Y) JHH7 cells stained for TOMM20. Boxed regions of interest (ROI #1 and ROI #2) are shown at higher magnification to the right. Scale bar, 25 µm. Images are representative of *n* = 3 independent biological replicates. **b**, Quantitative morphometric analysis of mitochondrial networks from the images in **a**, performed using the Mitochondria Analyzer plugin in Fiji/ImageJ, showing mean area, mean perimeter, mean aspect ratio, mean form factor, branches per mitochondrion, total branch length per mitochondrion, mean branch length, branch junctions per mitochondrion, branch end points per mitochondrion and mean branch diameter. Data are mean ± s.e.m. of *n* = 3 independent biological replicates, with at least 150 cells analyzed per condition; statistical significance was assessed by two-tailed unpaired Student’s *t*-test. **c**, Representative immunofluorescence images of lipid droplets (LD) stained with BODIPY 493/503 in parental and β-catenin (S33Y) cells. Two representative fields are shown for each condition, with the boxed region of Example 2 shown at higher magnification to the right. Scale bars, 25 µm. **d**, Quantification of lipid droplet number per cell from the images in **c**. Data are mean ± s.e.m. of *n* = 19 for parental and *n* = 26 for β-catenin (S33Y) cells, with at least 10 cells analyzed per field of view. Statistical significance was assessed on the experimental means by two-tailed unpaired Student’s *t*-test. Exact *p* values are shown in each panel.

Because mitochondrial network remodeling is often coupled to changes in lipid handling^36,37^, we next visualized lipid droplets by BODIPY 493/503 staining. β-catenin (S33Y) cells contained substantially fewer lipid droplets per cell when compared to parental cells (Fig. 3c,d). Consistent with a change in lipid handling, fatty acid metabolism was among the most positively enriched hallmark gene sets in β-catenin (S33Y) cells (Fig. 2a). Overall, these results demonstrate that activation of β-catenin promotes the formation of elongated, interconnected mitochondrial networks and decreases cellular lipid droplet content, raising the possibility that lipid stores are being mobilized to support mitochondrial metabolism.

### Active β-catenin changes lipidome composition and increases fatty acid oxidation

Two observations suggested that β-catenin activation alters cellular lipid handling: the reduction in lipid droplet content (Fig. 3c,d), and the transcriptional upregulation of genes related to fatty acid metabolism revealed by RNA sequencing (Fig. 2a). To more explicitly test this hypothesis, we performed untargeted lipidomics on parental and β-catenin (S33Y) cells. Principal component analysis separated the two genotypes along the first principal component, which accounted for 47.1% of the variance (Fig. 4a), indicating that β-catenin activation substantially remodels the cellular lipidome. We observed widespread changes across lipid classes, with species both increased and decreased in β-catenin (S33Y) cells (Fig. 4b). Grouping the differentially abundant species by pathway showed that triglyceride and diglyceride species associated with lipid droplets were broadly reduced, whereas acylcarnitines, which report on mitochondrial fatty acid oxidation, were increased (Fig. 4c). Sphingomyelin species were increased while ceramides were reduced, and membrane phospholipids showed species-specific changes in both directions (Fig. 4c).

**Fig. 4.**
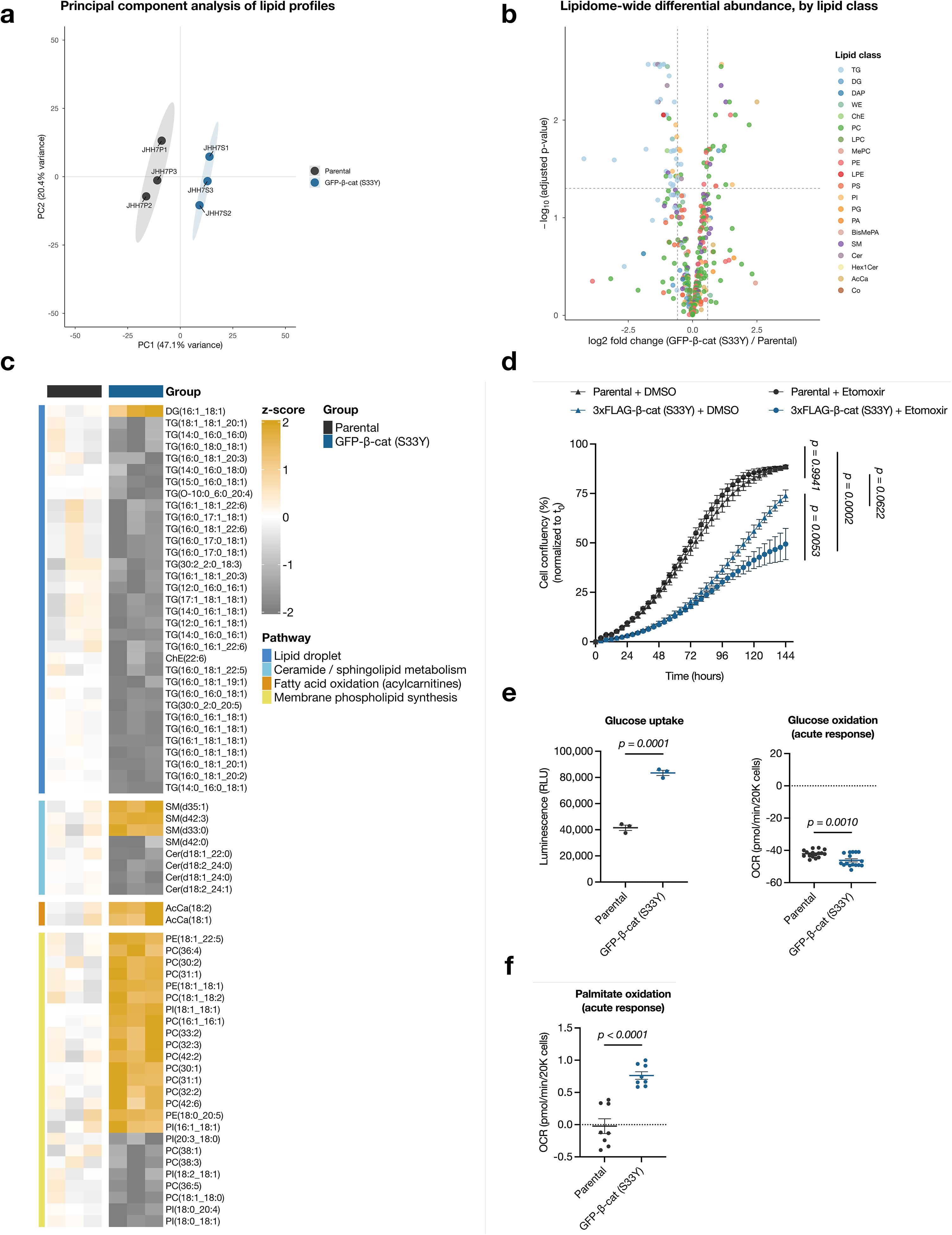
β-catenin activation remodels the cellular lipidome and shifts substrate oxidation toward fatty acids. **a**, Principal component analysis of lipid profiles from parental and β-catenin (S33Y) JHH7 cells, determined by untargeted lipidomics. Each point represents an independent biological replicate (*n* = 3 per group); ellipses indicate 95% confidence regions. **b**, Volcano plot of lipidome-wide differential abundance, showing log₂ fold change (β-catenin (S33Y)/parental) against −log₁₀ BH-adjusted *p* value. Points are colored by lipid class. Dashed lines indicate the significance thresholds: |log₂ fold change| = log₂(1.5) and BH-adjusted *p* = 0.05. **c**, Heatmap of differentially abundant lipid species in parental and β-catenin (S33Y) cells, shown as *z*-scores and grouped by pathway: lipid droplet, ceramide/sphingolipid metabolism, fatty acid oxidation (acylcarnitines) and membrane phospholipid synthesis. *n* = 3 independent biological replicates per group. **d**, Cell confluency over time for parental and β-catenin (S33Y) cells treated with vehicle (DMSO) or 20 µM etomoxir, measured by IncuCyte live-cell imaging every 4 h and normalized to t₀. Data are mean ± s.e.m. of *n* = 4 independent biological replicates; statistical significance was assessed at 144 h by two-way ANOVA with Fisher’s least significant difference test. **e**, Left, glucose uptake in parental and β-catenin (S33Y) cells, measured using the Glucose Uptake-Glo assay (*n* = 3 independent biological replicates per group), which detects intracellular 2-deoxyglucose-6-phosphate following incubation with 2-deoxyglucose, and expressed as relative luminescence units (RLU). Right, acute glucose oxidation, measured as the change in oxygen consumption rate (OCR) from baseline following acute injection of 25 mM glucose into substrate-deprived cells; OCR was normalized to 20,000 cells. Data are mean ± s.e.m. of *n* = 16 independent biological replicates; two-tailed unpaired Student’s *t*-test. **f**, Acute palmitate oxidation in parental and β-catenin (S33Y) cells, measured as the change in OCR from baseline following acute injection of 200 µM BSA-conjugated palmitate into substrate-deprived cells; OCR was normalized to 20,000 cells. Data are mean ± s.e.m. of *n* = 8 independent biological replicates; two-tailed unpaired Student’s *t*-test. Exact *p* values are shown in each panel.

The accumulation of acylcarnitines raised the possibility of altered flux through fatty acid oxidation. To test whether β-catenin (S33Y) cells depend on fatty acid oxidation for growth, we treated parental and β-catenin (S33Y) cells with the CPT1 inhibitor etomoxir and monitored confluence over time. Etomoxir had no effect on the growth of parental cells, whereas it significantly impaired the growth of β-catenin (S33Y) cells (Fig. 4d), indicating a selective dependence on fatty acid oxidation upon β-catenin activation.

We next examined how β-catenin activation affects the use of glucose and fatty acids as fuels for the TCA cycle and oxidative phosphorylation. To measure substrate oxidation directly, we deprived cells of exogenous substrates and then acutely provided either glucose or palmitate while monitoring oxygen consumption. β-catenin (S33Y) cells showed increased glucose uptake when compared to parental cells (Fig. 4e), yet their acute respiratory response to glucose was decreased relative to parental cells (Fig. 4e). Acute glucose exposure suppresses mitochondrial respiration in highly glycolytic cells, a phenomenon known as the Crabtree effect^38–40^, and the more pronounced suppression observed in β-catenin (S33Y) cells is consistent with reduced oxidation of glucose-derived carbon (Fig. 4e). In contrast, the acute respiratory response to palmitate was markedly increased in β-catenin (S33Y) cells (Fig. 4f), suggesting that β-catenin activation elicits a fuel preference for fatty acid substrates over carbohydrate substrates.

Overall, these results demonstrate that activation of β-catenin remodels the cellular lipidome and shifts oxidative substrate preference away from glucose and toward fatty acids.

### Hyperactivation of β-catenin rewires mitochondrial metabolism from carbohydrate oxidation to fatty acid oxidation

To determine directly which substrates supply carbon to the tricarboxylic acid (TCA) cycle, we performed stable isotope tracing in parental and β-catenin (S33Y) cells using uniformly labeled ¹³C substrates and measured fractional enrichment of TCA cycle intermediates. We used glucose and lactate as carbohydrate carbon sources, and palmitate and oleate as long-chain fatty acid carbon sources. Following incubation with [U-¹³C₆]-D-glucose, the M+2 labelling of α-ketoglutarate, succinate, fumarate and malate was reduced in β-catenin (S33Y) cells when compared to parental cells, whereas citrate labelling was unchanged (Fig. 5a). A similar pattern was observed with [U-¹³C₃]-L-lactate, which is converted to pyruvate by lactate dehydrogenase: M+2 labelling of α-ketoglutarate, succinate, fumarate and malate was reduced in β-catenin (S33Y) cells (Fig. 5b). Thus, the contribution of glucose- and lactate-derived carbon to the TCA cycle is diminished upon β-catenin activation. Consistent with a selective effect on carbohydrate-derived carbon, labelling from [U-¹³C₅]-L-glutamine was preserved, including modest increases in TCA intermediate labeling (Extended Data Fig. 3a).

**Fig. 5.**
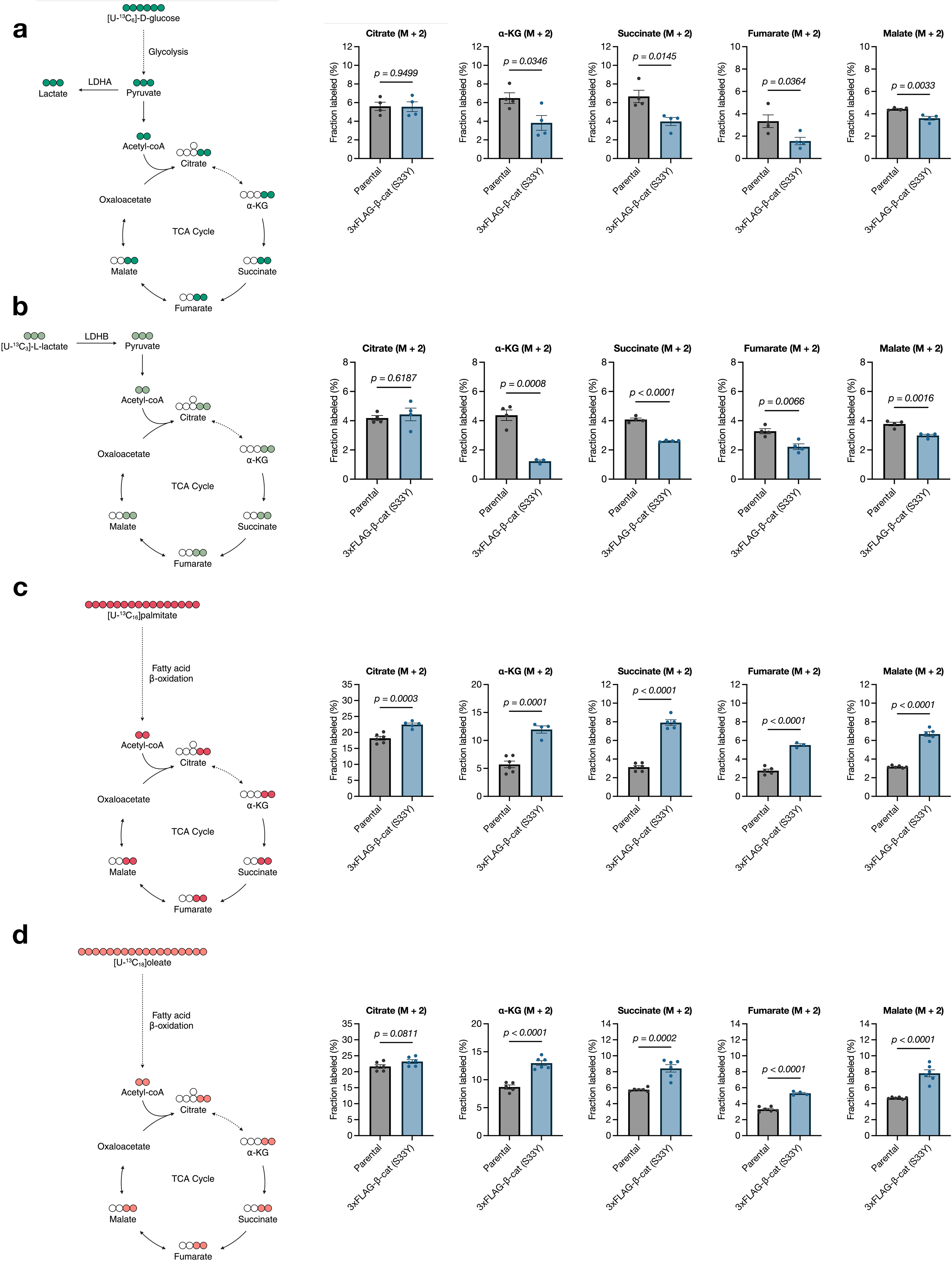
Stable isotope tracing reveals a shift from carbohydrate- to fatty acid-derived carbon entry into the TCA cycle. **a**, Left, schematic of [U-¹³C₆]-D-glucose metabolism through glycolysis and the TCA cycle; filled circles indicate ¹³C atoms. Right, fractional enrichment of M+2 citrate, α-ketoglutarate, succinate, fumarate and malate in parental and β-catenin (S33Y) JHH7 cells following 2 h incubation with [U-¹³C₆]-D-glucose. **b**, Left, schematic of [U-¹³C₃]-L-lactate metabolism, which enters the mitochondrion as pyruvate following conversion by lactate dehydrogenase B (LDHB). Right, fractional enrichment of the indicated M+2 TCA cycle intermediates following 1 h incubation with [U-¹³C₃]-L-lactate. **c**, Left, schematic of [U-¹³C₁₆]palmitate metabolism through fatty acid β-oxidation. Right, fractional enrichment of the indicated M+2 TCA cycle intermediates following 8 h incubation with [U-¹³C₁₆]palmitate in the presence of unlabeled glucose. **d**, Left, schematic of [U-¹³C₁₈]oleate metabolism through fatty acid β-oxidation. Right, fractional enrichment of the indicated M+2 TCA cycle intermediates following 8 h incubation with [U-¹³C₁₈]oleate in the presence of unlabeled glucose. Data are mean ± s.e.m. of *n* = 4 independent biological replicates for glucose and lactate tracing, and *n* = 6 independent biological replicates for palmitate and oleate tracing; statistical significance was assessed by two-tailed unpaired Student’s *t*-test. Exact *p* values are shown in each panel.

We next traced fatty acid-derived carbon. Following incubation with [U-¹³C₁₆]palmitate in the presence of unlabeled glucose, M+2 labelling of citrate, α-ketoglutarate, succinate, fumarate and malate was markedly increased in β-catenin (S33Y) cells (Fig. 5c). The same was observed with [U-¹³C₁₈]oleate (Fig. 5d). Because unlabeled glucose remained available throughout these experiments, the increased incorporation of fatty acid-derived carbon likely reflects a preference for fatty acid oxidation rather than a compensatory response to carbohydrate limitation.

Taken together, these results demonstrate that hyperactivation of β-catenin rewires mitochondrial substrate use, reducing the contribution of glucose- and lactate-derived carbon to the TCA cycle while increasing the oxidation of long-chain fatty acids.

## DISCUSSION

We set out to define the transcriptional regulators of metabolic state in liver cancer cells using the expression of *MPC1* as a marker gene. An unbiased CRISPR screen identified *CTNNB1* as a negative regulator of endogenous *MPC1* abundance, and pharmacologic or genetic activation of β-catenin reduced MPC1 and MPC2 at both the mRNA and protein level. Low MPC expression was also a feature of β-catenin-driven liver tumors in vivo. More importantly, β-catenin engineered a broader program that included reduced mitochondrial respiratory capacity, remodeled mitochondrial networks and alterations to the cellular lipidome. Functionally, β-catenin activation also shifted oxidative substrate use, with reduced contribution of glucose- and lactate-derived carbon to the TCA cycle and increased oxidation of palmitate and oleate. Our work is consistent with previous studies showing that β-catenin-activated hepatocellular carcinomas depend on fatty acid oxidation^27^.

Our results inform the mechanism(s) whereby β-catenin executes this fuel switch from carbohydrate to fatty acid-derived carbon substrates. Because the MPC is the only known carrier for pyruvate across the inner mitochondrial membrane^14–17^, its repression offers an explanation on that side of the switch: with reduced carrier abundance, pyruvate generated from glucose or lactate is less able to enter the mitochondrial matrix, while entry of fatty acid-derived acetyl-CoA, which does not require the MPC, is unaffected. *MPC1* repression is, however, one part of a broader program of gene expression that includes alterations to the mitochondrial respiratory chain, mitochondrial morphology and lipid content.

We have previously described transcriptional repression of the MPC in the mouse and human intestine, where low MPC expression supports stem cell function and is necessary and sufficient for tumor initiation^5–7^. Our results suggest that this MPC repression also operates in the liver, and may be a consequence of β-catenin hyperactivation, since β-catenin is one of the principal determinants of metabolic zonation^23,24^ and mutations affecting β-catenin are among the most frequent oncogenic events^26,27^. The broader context of the gene expression program, however, appear to differ between tissues. In the intestine, MPC loss is associated with a shift toward increased glucose uptake and glycolytic metabolism (accompanied by increased markers and functions of stemness), whereas in liver cancer cells we observe a shift toward fatty acid oxidation. Amongst other gene expression effects, this may also reflect the distinctive lipid-handling capacity of hepatocytes, and suggests that the metabolic outcome of MPC repression is determined by the context and tissue in which it occurs.

β-catenin activation also remodeled mitochondrial architecture and reduced lipid droplet number. In hepatocytes, mitochondrial network state and lipid droplet content are coupled through physical contacts between the two organelles, and this coupling is regulated by phosphorylation of PLIN5, with reduced contact site formation yielding fewer and smaller droplets^37^. Whether the elongated mitochondrial networks and depleted lipid droplet pools we observe reflect a change in mitochondria– lipid droplet coupling remains to be determined, but such a mechanism would provide a route by which stored lipid is directly delivered to mitochondria to support the increased fatty acid oxidation that we have measured.

A notable feature of our data is the disconnect between transcriptional and functional readouts of oxidative phosphorylation. β-catenin (S33Y) cells upregulated nuclear-encoded respiratory chain genes yet respired less, and the activities of the individual complexes were largely preserved. The defect instead correlated with the impaired assembly of Complexes III and IV and of Complex I-containing supercomplexes, together with reduced mitochondrial DNA copy number. Because Complexes I, III and IV contain mtDNA-encoded subunits whereas Complex II does not, the sparing of Complex II is consistent with a limitation imposed by reduced mitochondrial genome content. We interpret the transcriptional upregulation of the respiratory machinery as a compensatory response to this constraint, although the signals that couple mitochondrial genome content to nuclear gene expression in this setting remain to be determined.

We expect that MPC repression is only a part of this β-catenin driven metabolic program. The reduction in respiratory capacity, the changes in complex assembly and the loss of mitochondrial DNA are likely parallel outputs of β-catenin activation rather than downstream effects of MPC loss. We also note that a recent study reported that Wnt pathway activation does not reduce MPC levels and placed MPC loss upstream of Wnt signaling^41^. Those experiments were performed in fibroblasts and non-hepatic cell lines, and the relationship between Wnt signaling and the MPC may therefore be cell type-specific.

In summary, our results identify a mechanism by which an oncogenic signaling pathway engineers mitochondrial fuel selection. β-catenin was already known to determine the spatial organization of hepatic metabolism, with oxidative and gluconeogenic programs confined to the periportal zone and glycolytic and lipogenic programs to the perivenous zone where β-catenin is active^23,24^. The repression of the MPC provides one additional mechanism whereby this pathway can restrict entry of carbohydrate-derived carbon into the TCA cycle. Our data also raises the possibility that the same β-catenin-driven mechanism contributes to fuel selection in the normal liver as well as in tumors in which β-catenin is constitutively active. Given that HCC with hyperactive β-catenin constitutes a substantial fraction of human liver cancer and remains largely inaccessible to targeted therapy^26,27^, understanding how these tumors choose their substrates may help identify the metabolic dependencies that accompany that choice.

## METHODS

### Cell culture and treatment

Hep3B (HB-8064) and HEK293T (CRL-3216) cells were purchased from the American Type Culture Collection (ATCC). JHH7 (JCRB1031) and Huh-7 (JCRB0403) cells were obtained from the Japanese Collection of Research Bioresources (JCRB) Cell Bank. All cell lines were maintained in DMEM (Gibco, 10569-010) containing 25 mM glucose, 4 mM GlutaMAX and 1 mM sodium pyruvate, supplemented with 10% FBS (Sigma-Aldrich, F0926) and 50 U ml⁻¹ penicillin and 50 µg ml⁻¹ streptomycin (Gibco, 15140-122). Cells were maintained at 37 °C in a humidified incubator with 5% CO₂ and were used between passages 5 and 25. All cell lines were routinely tested for mycoplasma using the MycoAlert Mycoplasma Detection Kit (Lonza, LT07-318) and confirmed to be negative. Cell treatments were performed as indicated with DMSO (Hybri-Max; Sigma-Aldrich, D2650) as vehicle or with the following inhibitors: 20 µM CHIR99021 (Cayman Chemical, 13122) and 5 µM LY2090314 (Cayman Chemical, 22211).

### Generation of stable cell lines

Human CTNNB1 carrying the S33Y substitution was subcloned from pcDNA3-S33Y β-catenin^42^, a gift from Eric Fearon (Addgene plasmid #19286; RRID:Addgene_19286), into pLenti-CMV-Blast by restriction cloning, in frame with a GFP or 3xFLAG tag. Constructs were verified by Sanger sequencing. Second-generation lentiviral particles were produced in HEK293T cells by co-transfection of the transfer plasmid with psPAX2 (Addgene, 12260) and pMD2.G (Addgene, 12259) using Lipofectamine 2000 (Invitrogen, 11668019). Viral supernatants were collected at 24 and 48 h post-transfection, filtered through a 0.45 µm filter and applied to JHH7 cells at a 1:1 ratio with fresh culture medium in the presence of 10 µg ml⁻¹ polybrene (Sigma-Aldrich, TR-1003). Transduced cells were selected with 10 µg ml⁻¹ blasticidin (Gibco, A1113903). Expression of the tagged proteins was confirmed by immunoblotting, and β-catenin activation was confirmed by RT-qPCR. Parental JHH7 cells were used as controls throughout.

### SABER-FISH PER

Primer Exchange Reaction (PER) was carried out as described previously^43–45^. In brief, oligonucleotides were designed as described previously^46,47^ to ensure transcriptome-wide specificity. DNA oligonucleotides targeting RNAs of interest were appended with single-stranded DNA tails via catalytic PER hairpins. Sequences for each PER are listed in Supplementary Table 1.

### TRoUT-FISH screens

TRoUT-FISH screens were performed as previously described^28^ using the Human Epigenetic CRISPR Knockout Library, a gift from Kivanc Birsoy (Addgene plasmid #162256; RRID:Addgene_162256)^48^. Lentiviruses were produced in HEK293T cells by transfecting plasmids and packaging plasmids (psPAX2 and pMD2.G) with polyethylenimine transfection reagent. Media was collected every 24 hours for 96 hours total. Media was pooled, filtered with 0.45 μm PES filters, aliquoted, and frozen at −80 °C. The titer of lentivirus supernatant was determined by infecting target cells at various amounts of virus with polybrene followed by spinfection (30 °C for 60 minutes at 1,200 × g). The next day cells were reseeded and treated with puromycin. Viability was assessed 3 days post infection. Cells were infected at a multiplicity of infection (MOI) of approximately 0.4 via spinfection and treated with puromycin 24 hours post infection; puromycin selection persisted for 7 days. Cells were given fresh media for 6 hours followed by fixation with cold 4% PFA. Cells were permeabilized with PBS containing 0.1% Triton X-100 and Hoechst 33342 for 10 minutes rocking at 4 °C. Cells were incubated with 43 °C wHyb buffer^44,45^ for 30 minutes, then incubated overnight at 43 °C in hybridization buffer containing SABER-FISH PER purified DNA oligonucleotides. Subsequently, cells were washed two times with 43 °C wHyb c with 2× SSC buffer, and incubated at 30 °C for 1 hour in branch hybridization buffer containing SABER-FISH PER purified branch DNA oligonucleotides. After branching, cells were washed with 2× SSC and 37 °C PBS before incubation for 10 minutes in PBS containing fluorescently labeled secondary DNA oligonucleotides. After staining, cells were washed two times with 37 °C PBS and resuspended in 0.1% BSA (Millipore, 126609-5GM) in PBS for flow cytometry. Events were gated based on FSC and SSC to remove debris, then gated on DNA dye (violet laser) to ensure only singlets were sorted. A reference gene (EEF2) was used as a control to ensure cell quality and adequate staining. Cells were sorted based on fractions of the target gene distribution, sorting the lowest 10% and highest 10% of the total population. After sorting, cells were incubated in genomic DNA isolation buffer (0.5% SDS, 10 mM Tris pH 7.5, 10 mg ml⁻¹ proteinase K) and incubated at 56 °C for ≥36 hours with gentle agitation. Genomic isolation was performed with the Qiagen DNA Mini Kit (51304). Libraries were generated by PCR amplification and sequenced by 150 bp paired-end sequencing at the University of Utah High-Throughput Genomics Core Facility. Sequencing reads were quantified using BBTools Seal and analysis was performed by MAUDE using four bins representing deciles of the total population (bin A = 1–10, bin B = 11–20, bin C = 81–90, bin D = 91–100)^49^.

### Immunoblotting

Whole-cell extracts were prepared in 1× RIPA lysis buffer supplemented with 1 mM PMSF, cOmplete ULTRA EDTA-free protease inhibitor cocktail (Roche, 05892791001) and phosphatase inhibitor cocktail 2 (1:1,000; Sigma-Aldrich, P5726), and separated by 15% SDS-PAGE. Proteins were transferred to a nitrocellulose membrane (Cytiva, 10600002) and blocked with 5% nonfat dry milk. Immunoblotting was performed using the following primary antibodies, all from Cell Signaling Technology, at a 1:1,000 dilution: MPC1 (14462), MPC2 (46141), phospho-GSK3β (Ser9) (5558), GSK3β (9832), active β-catenin (19807), c-Myc (5605) and β-actin (8457). Membranes were incubated with the appropriate horseradish peroxidase-conjugated secondary antibodies at a 1:10,000 dilution, both from Bio-Rad Laboratories: Immun-Star goat anti-mouse (1705047) or Immun-Star goat anti-rabbit (1705046). Membranes were developed with Western Lightning Plus chemiluminescent detection solution (Revvity, NEL105001EA) and exposed to Blu-Lite UHC autoradiography film (MTC Bio, A8815). Densitometric analyses of immunoblots were performed using ImageJ as described in the figure legend.

### DNA extraction, RNA extraction and cDNA synthesis from cultured cells

For mitochondrial DNA (mtDNA) measurements, genomic DNA was extracted from cell pellets using the QIAamp DNA Blood Mini Kit (Qiagen). Total RNA was extracted from the indicated cultured cells using TRIzol reagent (Invitrogen) and Direct-zol RNA MiniPrep (Zymo Research) according to manufacturer’s protocol. 1–2 µg of RNA per sample was used for cDNA synthesis with the High-Capacity cDNA Reverse Transcription Kit (Applied Biosystems).

### RT-qPCR in cultured cells

For mtDNA measurements, 10–20 ng of genomic DNA were used. *D-Loop* and *CYTB* were used as mitochondrial probes and *ACTB* as nuclear DNA control. For the measurement of gene transcripts, 10– 20 ng of retrotranscribed cDNA were used and *RPLPO* was used as housekeeping gene for normalization purposes. In either case, real-time quantitative PCR was performed with PowerUp SYBR Green Master Mix for qPCR (Applied Biosystems) using the QuantStudio 7 Real-Time PCR system (Applied Biosystems). For each independent biological replicate, at least two technical replicates per gene and per sample were included. Sequences of used primers can be found in Supplementary Table 2. Data analysis was done according to the 2-ΔΔCT method. Statistical analysis was performed using One-way ANOVA with Dunnett correction or two-tailed unpaired Student’s *t*-test, and changes were considered significant if p-value < 0.05.

### RT-qPCR in mouse liver tissue

#### Animal husbandry

All procedures were approved by the Institutional Animal Care and Use Committee (IACUC) of the University of Utah. Mice were housed at 22–23 °C using a 12 h light/12 h dark cycle. Animals were maintained on a standard chow diet with ad libitum access (Envigo, Teklad 2020X). HCC tumor generation via hydrodynamic tail vein injection was performed in 6–8-week-old mice from C57BL/6J breeders purchased from The Jackson Laboratory. Equal numbers of male and female animals were used in experiments.

#### Hydrodynamic tail vein injection

HCC tumors and control liver tissue were generated in C57BL/6 mice by hydrodynamically injecting 10% of the mouse’s mass in mL of a plasmid–saline solution into the tail vein over 5–7 s. The plasmids pT3-N90-CTNNB1, pT3-EF1α-MYC-IRES-luciferase and pSB13, as well as the hydrodynamic tail vein injection protocols, were provided by Amaia Lujambio^50^. pSBtet-Pur was a gift from Eric Kowarz (Addgene plasmid #60507; RRID:Addgene_60507)^51^. HCC mice were injected with 1.425 µg ml⁻¹ pSB13, 5.7 µg ml⁻¹ pT3-EF1α-MYC-IRES-luciferase and 5 µg ml⁻¹ pT3-N90-CTNNB1. Control mice were injected with 1.425 µg ml⁻¹ pSB13 and 5.7 µg ml⁻¹ pSBtet-Pur. Mice were taken down between 4 and 6 weeks after the hydrodynamic tail vein injection, and the livers and tumors were flash frozen in liquid nitrogen.

#### RT-qPCR

Flash-frozen liver or tumor tissue (10–15 mg) was isolated in 2 mL Safe-Lock microcentrifuge tubes (Eppendorf) with a 5/16 in. diameter stainless steel ball (Grainger) chilled to −80 °C. Tissue was homogenized at 25 Hz for 30 s under liquid nitrogen using a CryoMill (Retsch). RNA was isolated using the RNeasy Mini Kit (Qiagen, 74104) according to the manufacturer’s specifications. cDNA was synthesized using qScript cDNA SuperMix (Quantabio) with 1 µg RNA as measured by NanoDrop One (Thermo Scientific). qPCR was carried out in a MicroAmp Optical 384-well reaction plate (Applied Biosystems) using PowerTrack SYBR Green Master Mix (Applied Biosystems) following the manufacturer’s instructions for a 10 µl reaction volume. *Actb* was used as the housekeeping gene, and the reaction was run on a QuantStudio 7 Flex (Applied Biosystems). Sequences of used primers can be found in Supplementary Table 2.

### RNA-seq

#### RNA-seq sample preparation and sequencing

Total RNA was isolated from parental and GFP-β-catenin (S33Y) JHH7 cells using TRIzol reagent (Invitrogen, 15596026) and the Direct-zol RNA MiniPrep kit (Zymo Research, R2052) according to the manufacturers’ protocols. RNA quantity and purity were assessed using a NanoDrop One spectrophotometer (Thermo Fisher Scientific), and RNA integrity was assessed using an Agilent RNA ScreenTape assay (5067-5576 and 5067-5577). Library preparation with poly(A) enrichment and sequencing were performed by Novogene using their human mRNA sequencing service on a NovaSeq X Plus system (Illumina) with 150 bp paired-end reads.

#### RNA-seq differential expression analysis

Raw counts were filtered to remove genes with fewer than 10 total reads across samples, and differential expression analysis was performed using DESeq2 (v.1.50.2) in R (v.4.5.2), modeling expression as a function of experimental group (GFP-β-catenin (S33Y) versus parental), with parental set as the reference level. Genes with adjusted *p* < 0.05 were considered differentially expressed. For visualization of individual genes, per-sample log₂ fold change values were calculated by log₂-transforming DESeq2-normalized counts (+1 pseudocount) and subtracting, for each gene, the mean log₂-transformed value across parental samples. Heatmaps of ETC gene expression were generated from *z*-scored, DESeq2-normalized counts using pheatmap (v.1.0.13), with genes manually ordered by respiratory complex.

#### Gene set enrichment analysis (GSEA)

GSEA was performed using genes ranked by Wald statistic from the DESeq2 model, using the clusterProfiler package (v.4.18.4) in R (v.4.5.2). Enrichment was assessed against MSigDB Hallmark gene sets (v.26.1.0), with pathways considered significant at adjusted *p* < 0.01. Enrichment was also assessed against mitochondrial pathway gene sets curated from MitoCarta 3.0^52^, with redundant subcategories (individual complex subunit and assembly factor listings already represented within their parent complex categories) removed prior to analysis. Minimum and maximum gene set sizes were set to 10 and 500, respectively, and pathways were considered significant at adjusted *p* < 0.01.

### Seahorse extracellular flux analysis

Assays were performed on a Seahorse XF Pro Analyzer (Agilent) using the Seahorse XF Cell Mito Stress Test Kit (Agilent, 103015-100). Parental or GFP-β-catenin (S33Y) JHH7 cells were plated on a 96-well Seahorse plate at a density of 20,000 cells per well in complete DMEM containing 10% FBS and penicillin–streptomycin, and allowed to attach overnight. Following each assay, cells were stained with 5 µg ml⁻¹ Hoechst 33342 (Invitrogen, H3570) and imaged on a BioTek Cytation 1 Cell Imaging Multimode Reader (Agilent) for cell counting, and oxygen consumption rates were normalized to cell number. Data were analyzed in Seahorse Wave software (Agilent).

#### Mitochondrial stress test

On the day of the experiment, the medium was replaced with Seahorse XF DMEM medium (pH 7.4) supplemented with 10 mM glucose, 1 mM sodium pyruvate and 2 mM L-glutamine. The assay was performed using final concentrations of 1.5 µM oligomycin, 1 µM FCCP and 0.5 µM rotenone plus 0.5 µM antimycin A, with the standard protocol consisting of three measurement cycles for each assay phase (basal, oligomycin, FCCP and rotenone/antimycin A), each comprising 3 min mixing, 0 min waiting and 3 min measuring. Data were analyzed using the XF Mitochondrial Stress Test Report Generator.

#### Substrate oxidation stress test

On the day of the experiment, the medium was replaced with Seahorse XF DMEM medium (pH 7.4) without glucose, sodium pyruvate or L-glutamine, and cells were starved for 1 h at 37 °C. The assay was performed by acutely injecting 25 mM glucose (Gibco, A24940-01) or 200 µM BSA-conjugated palmitate (Cayman Chemical, 29558), followed by final concentrations of 1.5 µM oligomycin, 1 µM FCCP and 0.5 µM rotenone plus 0.5 µM antimycin A. The protocol consisted of three measurement cycles for each assay phase, with five measurement cycles following the acute injection, each comprising 3 min mixing, 0 min waiting and 3 min measuring. Data were analyzed using the XF Mitochondrial Stress Test Acute Response Report Generator. The acute response was calculated as the difference between the OCR after substrate injection and the baseline OCR immediately before injection.

### Crude mitochondrial isolation for SDS-PAGE and BN-PAGE

Crude mitochondrial isolations were prepared as previously described^9,53^. Parental or 3xFLAG-β-catenin (S33Y) JHH7 cells were grown to full confluency, collected by trypsinization and centrifuged at 1,200 rpm for 10 min at 25 °C. The pellet was washed with PBS and frozen at −80 °C. The pellet was resuspended in 1 ml of CP-1 buffer (100 mM KCl, 50 mM Tris-HCl, 2 mM EGTA, pH 7.4), homogenized five times by passage through a 21G needle, and centrifuged at 700 × *g* for 10 min at 4 °C. The supernatant was transferred to a new tube and centrifuged at 10,500 × *g* for 10 min at 4 °C. The pellet, containing crude mitochondria, was washed twice with 1 ml of CP-1 buffer and resuspended in 50 µl of CP-1 buffer. Protein concentration was measured and normalized using Quick Start Bradford 1× Dye Reagent (Bio-Rad Laboratories, 5000205).

#### SDS-PAGE and immunoblotting of mitochondrial lysates

Crude mitochondrial preparations were resuspended in 6× SDS sample buffer, normalized to total protein content and separated by 15% SDS-PAGE. Mitochondrial proteins were transferred to a nitrocellulose membrane (Cytiva, 10600002) and blocked with 5% nonfat dry milk. Immunoblotting was performed using the following primary antibodies at a 1:1,000 dilution: NDUFS2 (Abcam, ab110249), SDHA (Abcam, ab14715), UQCRC2 (Abcam, ab14745), COXIV (Proteintech, 11242-1-AP), ATP5a (Abcam, ab14748) and Hsp60 (Novus, NBP2-12734). Secondary antibody incubation and detection were performed as described above.

#### Blue native PAGE (BN-PAGE) and immunoblotting of mitochondrial lysates

Crude mitochondrial preparations were normalized to total protein content and separated by 3–12% BN-PAGE (Invitrogen, BN2011BX10) as previously described^9,53^. Briefly, 100 µg of mitochondria was lysed on ice in 100 µl of 1× pink NativePAGE sample buffer (Invitrogen, BN2003) containing 1% digitonin (GoldBio, D-180) for 15 min and centrifuged at 20,000 × *g* for 20 min at 4 °C. Cleared mitochondrial lysate was transferred to a new 1.5 ml Eppendorf tube and 5 µl of NativePAGE 5% G-250 sample additive (Invitrogen, BN2004) was added. Mitochondrial protein extracts (10 µg) were separated by 3–12% BN-PAGE using the NativePAGE system (Invitrogen, BN2007). Mitochondrial proteins were transferred to a polyvinylidene difluoride membrane (MilliporeSigma, IPVH00010) and blocked with 5% nonfat dry milk. Immunoblotting was performed using the following primary antibodies at a 1:1,000 dilution: NDUFS2 (Abcam, ab110249), SDHA (Abcam, ab14715), UQCRC2 (Abcam, ab14745), COXIV (Proteintech, 11242-1-AP) and ATP5a (Abcam, ab14748). Secondary antibody incubation and detection were performed as described above.

### Mitochondrial complex activity assays

For enzymatic activity assays, mitochondria were isolated as described above and solubilized in CP-1 buffer containing 1% (w/v) digitonin at 1 mg protein ml⁻¹ immediately before use. The enzymatic activities of mitochondrial respiratory chain complexes I, II, III, IV and V were measured spectrophotometrically in digitonin-solubilized mitochondria isolated from JHH7 cells stably expressing 3xFLAG-β-catenin (S33Y) or parental JHH7 cells, adapting established protocols^54,55^ to a 96-well microplate format. All assays were performed in a final reaction volume of 200 µl on a BioTek Synergy Neo2 multi-mode microplate reader, monitoring absorbance kinetically at 37 °C. Citrate synthase activity was additionally measured in every sample as a marker of mitochondrial matrix content and used to normalize the activity of each respiratory chain complex. For each complex, a parallel reaction containing a complex-specific inhibitor was run alongside the uninhibited reaction, and complex-specific activity was calculated as the inhibitor-sensitive rate (uninhibited rate minus inhibited rate). Activities were calculated from the linear rate of absorbance change as: activity (nmol min⁻¹ mg⁻¹ protein) = (ΔA/min × 0.2 ml × 1,000) / (ε [mM⁻¹ cm⁻¹] × protein [mg] × path length [cm]), assuming a 1 cm path length, and expressed relative to citrate synthase activity and to the mean value of parental controls.

#### Complex I activity assay

Complex I (NADH:ubiquinone oxidoreductase) activity was measured as the rotenone-sensitive rate of NADH oxidation at 340 nm (ε = 6.22 mM⁻¹ cm⁻¹). Reactions contained 5 µg mitochondrial protein, 50 mM potassium phosphate (pH 7.5), 3 mg ml⁻¹ BSA, 300 µM KCN, 100 µM NADH and either vehicle (ethanol) or 10 µM rotenone. Baseline absorbance was recorded for 2 min, the reaction was initiated with 60 µM ubiquinone-1, and absorbance was recorded for a further 5 min.

#### Complex II activity assay

Complex II (succinate dehydrogenase) activity was measured as the malonate-sensitive rate of 2,6-dichlorophenolindophenol (DCPIP) reduction at 600 nm (ε = 19.1 mM⁻¹ cm⁻¹). Reactions contained 4 µg mitochondrial protein, 25 mM potassium phosphate (pH 7.5), 1 mg ml⁻¹ BSA, 20 mM succinate, 300 µM KCN and 75 µM DCPIP, and were pre-incubated at 37 °C for 10 min to activate succinate dehydrogenase. Baseline absorbance was recorded for 2 min, the reaction was initiated with 50 µM decylubiquinone, and absorbance was recorded for 3 min; 10 mM malonate was then added and absorbance recorded for a further 2 min to define the malonate-sensitive rate.

#### Complex III activity assay

Complex III (ubiquinol:cytochrome *c* oxidoreductase) activity was measured as the antimycin A-sensitive rate of cytochrome *c* reduction at 550 nm (ε = 18.5 mM⁻¹ cm⁻¹). Reactions contained 5 µg mitochondrial protein, 25 mM potassium phosphate (pH 7.5), 1 mg ml⁻¹ BSA, 500 µM KCN, 75 µM oxidized cytochrome *c* and either vehicle (ethanol) or 10 µg ml⁻¹ antimycin A. Baseline absorbance was recorded for 2 min, the reaction was initiated with 10 µM decylubiquinol, and absorbance was recorded for 5 min.

#### Complex I and III activity assay

Complex I and III (NADH:cytochrome *c* oxidoreductase) activity was measured as the rotenone-sensitive rate of cytochrome *c* reduction at 550 nm (ε = 18.5 mM⁻¹ cm⁻¹). Reactions contained 5 µg mitochondrial protein, 50 mM potassium phosphate (pH 7.5), 1 mg ml⁻¹ BSA, 300 µM KCN, 50 µM oxidized cytochrome *c* and either vehicle (ethanol) or 10 µM rotenone. Baseline absorbance was recorded for 2 min, the reaction was initiated with 200 µM NADH, and absorbance was recorded for 5 min.

#### Complex IV activity assay

Complex IV (cytochrome *c* oxidase) activity was measured as the KCN-sensitive rate of reduced cytochrome *c* oxidation at 550 nm (ε = 18.5 mM⁻¹ cm⁻¹). Reactions contained 25 mM potassium phosphate (pH 7.0) and 50 µM reduced cytochrome *c*. Baseline absorbance was recorded for 2 min, the reaction was initiated by addition of 5 µg mitochondrial protein, and absorbance was recorded for 5 min; 300 µM KCN was then added and absorbance recorded for 2 min to define the KCN-sensitive rate.

#### Complex V activity assay

Complex V (F₁F₀-ATP synthase) activity was measured as the oligomycin-sensitive rate of ATP hydrolysis, coupled to NADH oxidation through pyruvate kinase and lactate dehydrogenase and monitored at 340 nm (ε = 6.22 mM⁻¹ cm⁻¹). Reactions contained 5 µg mitochondrial protein, 50 mM Tris-HCl (pH 8.0), 5 mg ml⁻¹ BSA, 20 mM MgCl₂, 50 mM KCl, 2.5 mM ATP, 10 mM phosphoenolpyruvate, 15 µM CCCP (to dissipate the membrane potential and prevent reverse electron transport from confounding ATP hydrolysis), 5 µM antimycin A (to block respiratory chain electron transport) and excess coupling enzymes (pyruvate kinase and lactate dehydrogenase). Mitochondria were pre-incubated at 37 °C for 1 min before addition of assay buffer, and baseline absorbance was recorded for 5 min. The reaction was initiated with NADH (200 µM final), and absorbance was recorded for 3 min; 10 µM oligomycin was then added and absorbance recorded for 2 min to define the oligomycin-sensitive rate.

### Mitochondrial membrane potential

Mitochondrial membrane potential was assessed in JHH7 cells stably expressing 3xFLAG-β-catenin (S33Y) or parental JHH7 cells as the ratio of MitoTracker Red CMXRos to MitoTracker Green FM fluorescence, which normalizes membrane-potential-dependent dye accumulation to mitochondrial mass^8^. Briefly, 300,000 cells per well were incubated with 20 nM MitoTracker Red CMXRos (Invitrogen, M7512) and 20 nM MitoTracker Green FM (Invitrogen, M7514) for 20 min at 37 °C. As a positive control for membrane depolarization, a parallel set of cells was treated with 25 µM CCCP for 2 h to dissipate mitochondrial membrane potential. Cells were then washed with PBS, trypsinized and resuspended in PBS containing 1 mM EDTA and 1 µg ml⁻¹ DAPI for flow cytometry. Samples were acquired on a CytoFLEX LXR flow cytometer (Beckman Coulter). Cells were gated sequentially on scatter (FSC-A versus SSC-A), singlets (FSC-A versus FSC-width) and DAPI-negative (live) populations using unstained controls. Mitochondrial membrane potential was calculated for each sample as the ratio of mean MitoTracker Red to mean MitoTracker Green fluorescence intensity within the live-gated population.

### Immunofluorescence

For TOMM20 immunostaining, parental or 3xFLAG-β-catenin (S33Y) JHH7 cells were plated on glass coverslips with poly-L-lysine coating (Neuvitro, GG-12-1.5-PLL). Cells were fixed for 15 min with 4% paraformaldehyde in PBS and permeabilized for 7 min with 0.1% Triton X-100 in PBS. Fixed samples were washed four times with PBS and blocked overnight with 5% bovine serum albumin (BSA) in PBS at 4 °C. Immunofluorescence was performed with TOMM20 primary antibody at a 1:200 dilution (Proteintech, 11802-1-AP) at room temperature for 1 h. Coverslips were washed four times with PBS and incubated at room temperature for 1 h with Alexa Fluor 568 goat anti-rabbit secondary antibody (Life Technologies, A11011) at a 1:200 dilution. After incubation, the coverslips were washed four times with PBS and mounted in ProLong Diamond Antifade mountant with DAPI (Invitrogen, P36962). To label lipid droplets, BODIPY 493/503 (Life Technologies, D3922) was added to live cells at 200 ng ml⁻¹ immediately before imaging and was present during imaging. Images were acquired with a Zeiss LSM 880 confocal microscope with Airyscan and are shown as maximum projections. Quantitative analysis of mitochondrial morphology was performed using the Mitochondria Analyzer plugin in Fiji/ImageJ^56^. Lipid droplet number was quantified using the Analyze Particles function in ImageJ on thresholded images, with size settings from 0.1 to 100 square pixels and circularity from 0 to 1, as previously described^36^.

### Lipid extraction

Parental or GFP-β-catenin (S33Y) JHH7 cells were plated on 6 cm plates and grown for 2 days in complete DMEM (Gibco, 10569-010) with 10% dialyzed FBS (Gibco, 26400-044) to approximately 90% confluency. Medium was refreshed 2 h before extraction. To extract, cells were washed twice with cold PBS, scraped into additional PBS using a cell lifter (CellTreat, 229305), and transferred to a 1.5 ml microcentrifuge tube. After centrifugation, PBS was removed and cells were lysed in 1 ml extraction solution containing 75% HPLC-grade methyl tert-butyl ether (MTBE), 24% HPLC-grade methanol and 1% SPLASH LIPIDOMIX Mass Spec Standard (Avanti Polar Lipids, 330707). After 15 min of incubation on ice with intermittent vortexing, 200 µl HPLC-grade water was added, and the lysates were vortexed and incubated on ice for an additional 5 min. After centrifugation, the top organic layer was transferred to a glass vial and dried under gaseous nitrogen. The lipid extracts were resuspended in 200 µl of 2:1:1 HPLC-grade isopropyl alcohol, acetonitrile and water, and transferred to amber mass spectrometry vials (Agilent, 5182-0716) with glass inserts.

### LC–MS/MS lipidomics

The lipid extracts were analyzed by LC–MS/MS using a Vanquish HPLC system (Thermo Fisher Scientific) and a Q Exactive HF Orbitrap mass spectrometer (Thermo Fisher Scientific). Lipid separation was performed on an Acquity UPLC CSH C18 column (2.1 mm × 100 mm, 1.7 µm particle size, 130 Å pore size; Waters, 186005297) with a constant flow rate of 350 µl min⁻¹ and a chromatography gradient of: 0 min, 30% B; 5 min, 43% B; 5.1 min, 50% B; 14 min, 70% B; 21 min, 99% B; 24 min, 99% B; 24.1 min, 30% B; 31 min, 30% B. Solvent A was 10 mM ammonium formate and 0.1% formic acid in acetonitrile:water (60:40), and solvent B was 10 mM ammonium formate and 0.1% formic acid in isopropyl alcohol:acetonitrile:water (90:9:1). Autosampler temperature was 4 °C, column temperature was 30 °C and injection volume was 2 µl. Samples were injected into the mass spectrometer by electrospray ionization operating in positive ion mode and analyzed using a ddMS2 method with a resolving power of 35,000 at *m/z* 200. Peaks were identified using LipidSearch 5.0 (Thermo Fisher Scientific).

### Lipidomics data analysis

Lipidomic profiling was performed on JHH7 cells stably expressing GFP-β-catenin (S33Y). Peak-integrated lipid intensities were used as the starting point for all downstream bioinformatic analysis, which was performed in R (v.4.5.0) using the packages ggplot2 (v.4.0.3), dplyr (v.1.2.1), tidyr (v.1.3.2), tibble (v.3.3.1) and purrr (v.1.2.2) from the tidyverse, limma (v.3.66.0), ComplexHeatmap (v.2.26.1), circlize (v.0.4.18), ggrepel (v.0.9.8), ggpubr (v.1.0.0) and RColorBrewer (v.1.1-3).

Analytical reproducibility was assessed as the relative standard deviation (RSD = SD/mean × 100) of raw, non-imputed peak intensities, computed separately across the three biological replicates per genotype and across two pooled quality-control (QC) injections. Deuterated internal standards were used for normalization, and endogenous lipid species were required to be detected in at least 2 of 3 biological replicates. Normalized intensities were log₂-transformed for all downstream analyses. This pipeline retained 384 lipid species across 20 lipid classes.

Principal component analysis was performed on autoscaled (mean-centered, unit-variance-scaled) log₂ normalized intensities across all 384 species (prcomp, R). Differential lipid abundance between genotypes was assessed by moderated limma *t*-test with empirical Bayes variance shrinkage, and *p* values were adjusted for multiple testing by Benjamini–Hochberg (BH). Lipid species were called significant at BH-adjusted *p* < 0.05 and |log₂ fold change| > log₂(1.5). Volcano plots display log₂ fold change (*x* axis) against −log₁₀(BH-adjusted *p* value) (*y* axis); lipid-class volcano plots color each point by its lipid class.

Heatmaps were generated with ComplexHeatmap. For each significant lipid species (BH-adjusted *p* < 0.05, |log₂FC| > log₂(1.5)), row-wise *z*-scores were computed by centering on the mean of the control-group replicates and scaling by the standard deviation across all replicates. Rows were hierarchically clustered (Euclidean distance, complete linkage) within four curated pathway groups (lipid droplet lipids: triacylglycerol, diacylglycerol, cholesteryl ester; sphingolipid metabolism: ceramide, hexosylceramide, sphingomyelin; fatty acid oxidation: acylcarnitine; membrane phospholipid synthesis: phosphatidylcholine, phosphatidylethanolamine, phosphatidylserine, phosphatidylinositol); columns were grouped by genotype without clustering.

The total acyl-chain carbon and double-bond count of each lipid species was parsed from its annotated composition string and classified as saturated (0 double bonds), monounsaturated (1 double bond) or polyunsaturated (≥2 double bonds). Each category’s contribution was calculated as the abundance-weighted share of total normalized signal per genotype, both across the whole lipidome and within individual lipid classes. Per-sample saturation composition was compared between genotypes by two-sided *t*-test.

### Cell proliferation assays

Parental or 3xFLAG-β-catenin (S33Y) JHH7 cells were seeded in 24-well plates at a density of 20,000 cells per well in glucose-free DMEM containing 4 mM L-glutamine (Gibco, 11966-025), supplemented with 25 mM glucose (Gibco, A24940-01) and 10% charcoal-stripped FBS (Gibco, A33821-01), with either DMSO as vehicle or 20 µM etomoxir (Cayman Chemical, 11969). Plates were imaged every 4 h for 144 h on an IncuCyte S3 live-cell imaging system (Sartorius) at 37 °C and 5% CO₂, with four images acquired per well using a 10× objective. Phase-contrast confluency was calculated using IncuCyte S3 software (v.2025B, Sartorius) and expressed as percentage confluency normalized to t₀.

### Glucose uptake assay

Parental or GFP-β-catenin (S33Y) JHH7 cells were seeded in 96-well plates at a density of 20,000 cells per well the day before the experiment. For glucose uptake, cells were starved of glucose for 1 h before measurement using the Glucose Uptake-Glo Assay (Promega, J1341) according to the manufacturer’s instructions. Luminescence was recorded on a BioTek Synergy Neo2 multi-mode microplate reader and normalized to 20,000 cells per well.

### Stable isotope tracing

Parental or 3xFLAG-β-catenin (S33Y) JHH7 cells were seeded in 6-well plates at a density of 0.5 × 10⁶ cells per well and grown for 2 days to 70–80% confluency before tracing. All labeled substrates were from Cambridge Isotope Laboratories.

#### Glucose and lactate tracing

Cells were washed and incubated for 1 h in DMEM lacking glucose, glutamine and pyruvate (Gibco, A14430-01). Cells were then labeled in glucose- and pyruvate-free DMEM containing 4 mM L-glutamine (Gibco, 11966-025) supplemented with 10% dialyzed FBS (Gibco, 26400-044) and either 25 mM [U-¹³C₆]-D-glucose (CLM-1396-10) for 2 h or 1.6 mM [U-¹³C₃]-L-lactate (CLM-1579-0.5) for 1 h.

#### Glutamine tracing

Cells were washed and incubated for 1 h in DMEM lacking glucose, glutamine and pyruvate (Gibco, A14430-01). Cells were then labeled for 2 h in the same medium supplemented with 25 mM glucose, 10% dialyzed FBS (Gibco, 26400-044) and 2 mM [U-¹³C₅]-L-glutamine (CLM-1822-H-0.1).

#### Palmitate and oleate tracing

Cells were washed and incubated for 1–2 h in HBSS (Gibco, 14025-076). Cells were then labeled for 8 h in glucose- and pyruvate-free DMEM containing 4 mM L-glutamine (Gibco, 11966-025) supplemented with 25 mM unlabeled glucose, 10% charcoal-stripped FBS (Gibco, A33821-01) and 200 µM [U-¹³C₁₆]palmitate (CLM-409) or [U-¹³C₁₈]oleate (CLM-460), conjugated to 2% fatty acid-free BSA.

#### Sample collection

At the end of the labeling period, cells were washed once with cold 1× PBS, and cells and media were collected and extracted as described below.

### Metabolite extraction and data analysis

Cell and media samples were extracted in −80 °C 80:20 methanol:water, vortexed, incubated on dry ice for 10 min and centrifuged at 16,000 × *g* for 25 min, with the supernatant used for LC–MS analysis. Extracts were analyzed within 24 h by liquid chromatography coupled to mass spectrometry (LC–MS). The LC–MS method was based on hydrophilic interaction chromatography (HILIC) coupled to an Orbitrap Exploris 240 mass spectrometer (Thermo Scientific)^57^. LC separation was performed on an XBridge BEH Amide column (2.1 × 150 mm, 3.5 µm particle size; Waters). Solvent A was H₂O:acetonitrile (95:5) with 20 mM ammonium acetate and 20 mM ammonium hydroxide, and solvent B was acetonitrile:H₂O (90:10) with 20 mM ammonium acetate and 20 mM ammonium hydroxide. The gradient was 0 min, 90% B; 2 min, 90% B; 3 min, 75% B; 5 min, 75% B; 6 min, 75% B; 7 min, 75% B; 8 min, 70% B; 9 min, 70% B; 10 min, 50% B; 12 min, 50% B; 13 min, 25% B; 14 min, 25% B; 16 min, 0% B; 18 min, 0% B; 20 min, 0% B; 21 min, 90% B; 25 min, 90% B. The following parameters were maintained during LC analysis: flow rate 150 µl min⁻¹, column temperature 25 °C, injection volume 5 µl and autosampler temperature 5 °C. For the detection of metabolites, the mass spectrometer was operated in both negative and positive ion mode. The following parameters were maintained during MS analysis: resolution of 180,000 at *m/z* 200, automatic gain control (AGC) target at 3 × 10⁶, maximum injection time of 30 ms and scan range of *m/z* 70–1,000. Raw LC–MS data were converted to mzXML format using the command-line msconvert utility^58^. Data were analyzed with El-MAVEN (v.12) and all isotope labeling patterns were corrected for natural ¹³C abundance using AccuCor^59^.

### Quantification and statistical analysis

Quantification and statistical analyses were performed as described in the figure legends. Comparisons between two groups were analyzed using a two-tailed unpaired Student’s *t*-test. Comparisons between three or more groups were analyzed using one-way analysis of variance (ANOVA) followed by Dunnett’s multiple comparisons test, or two-way ANOVA followed by Fisher’s least significant difference (LSD) test. Unless otherwise stated, analyses were performed on at least three independent biological replicates using GraphPad Prism (v.11.1.0; GraphPad Software). Data are shown as mean ± s.e.m., and *p* < 0.05 was considered statistically significant.

## DATA AVAILABILITY

We included all the data necessary in the interpretation of this study. Reagents and constructs are available upon request.

## ACKNOWLDEGEMENTS

We wish to acknowledge the support of the Metabolic Phenotyping Core and the Flow Cytometry Core Facilities at the Health Sciences Center at the University of Utah, and we thank Ying Li and Xue Yin, and James Marvin and Sreeja Govindarajan, respectively, for their technical support. We also acknowledge the Huck Institutes’ Metabolomics Core Facility (RRID:SCR_023864) at The Pennsylvania State University for use of the OE 240 LCMS, and Sergei Koshkin for helpful discussions on sample preparation and analysis. We thank members of the Rutter laboratory for their helpful feedback and discussions. All schematics were created in BioRender.

## AUTHOR CONTRIBUTIONS

L.C.-R. and J.R. conceived the study. L.C.-R. designed and performed the majority of the experiments and analyzed the data. Q.A.A. performed and analyzed the mitochondrial membrane potential and complex activity assays, and analyzed the lipidomics data, with assistance from M.-Y.J. on the complex activity analysis. N.A.S. and P.P. performed all analyses of the stable isotope tracing studies. J.W. performed RT-qPCR on mouse tissue, ran the lipidomics, and, with J.N.V.-S., provided guidance on the tracing studies. G.L. and S.F. performed cloning of the GFP- and 3xFLAG-tagged β-catenin constructs. N.A. and J.C.Y. performed immunoblotting and RT-qPCR, respectively, of MPC expression following CHIR99021 and LY2090314 treatment. Á.J.N.-P. and L.G.M. performed RT-qPCR analysis of parental versus β-catenin (S33Y) cells, and Á.J.N.-P. performed RT-qPCR analysis of mitochondrial DNA content. J.F. performed mitochondrial isolation for BN-PAGE and SDS-PAGE. K.H. performed the RNA sequencing data analysis. A.J.B. performed the TRoUT-FISH screen. M.A.M. performed the lipid droplet image analysis. G.S.D. provided liver tissue from the Sleeping Beauty model of hepatocellular carcinoma. G.S.D., J.L.-S. and M.R.M. provided additional study guidance. L.C.-R., G.S.D., J.L.-S., M.R.M. and J.R. provided funding support. J.R. supervised the project. L.C.-R. wrote the original draft, and L.C.-R. and J.R. revised the manuscript with input from all authors.

## ABBREVIATIONS

BN-PAGE: blue native polyacrylamide gel electrophoresis
BSA: bovine serum albumin
CPT1: carnitine palmitoyltransferase 1
DMSO: dimethyl sulfoxide
FACS: fluorescence-activated cell sorting
FAO: fatty acid oxidation
FCCP: carbonyl cyanide-4-(trifluoromethoxy)phenylhydrazone
GSEA: gene set enrichment analysis
HCC: hepatocellular carcinoma
LD: lipid droplet
MAUDE: mean alterations using discrete expression
MMP: mitochondrial membrane potential
MPC: mitochondrial pyruvate carrier
mtDNA: mitochondrial DNA
NES: normalized enrichment score
OCR: oxygen consumption rate
OXPHOS: oxidative phosphorylation
PCA: principal component analysis
RLU: relative luminescence units
ROI: region of interest
RRA: robust rank aggregation
RT-qPCR: reverse transcription quantitative polymerase chain reaction
SDS-PAGE: sodium dodecyl sulfate polyacrylamide gel electrophoresis
TCA: tricarboxylic acid;
TRoUT-FISH: Targeted Readout to Understand Transcription via Fluorescent In Situ Hybridization

---

**Extended Data Fig. 1.**
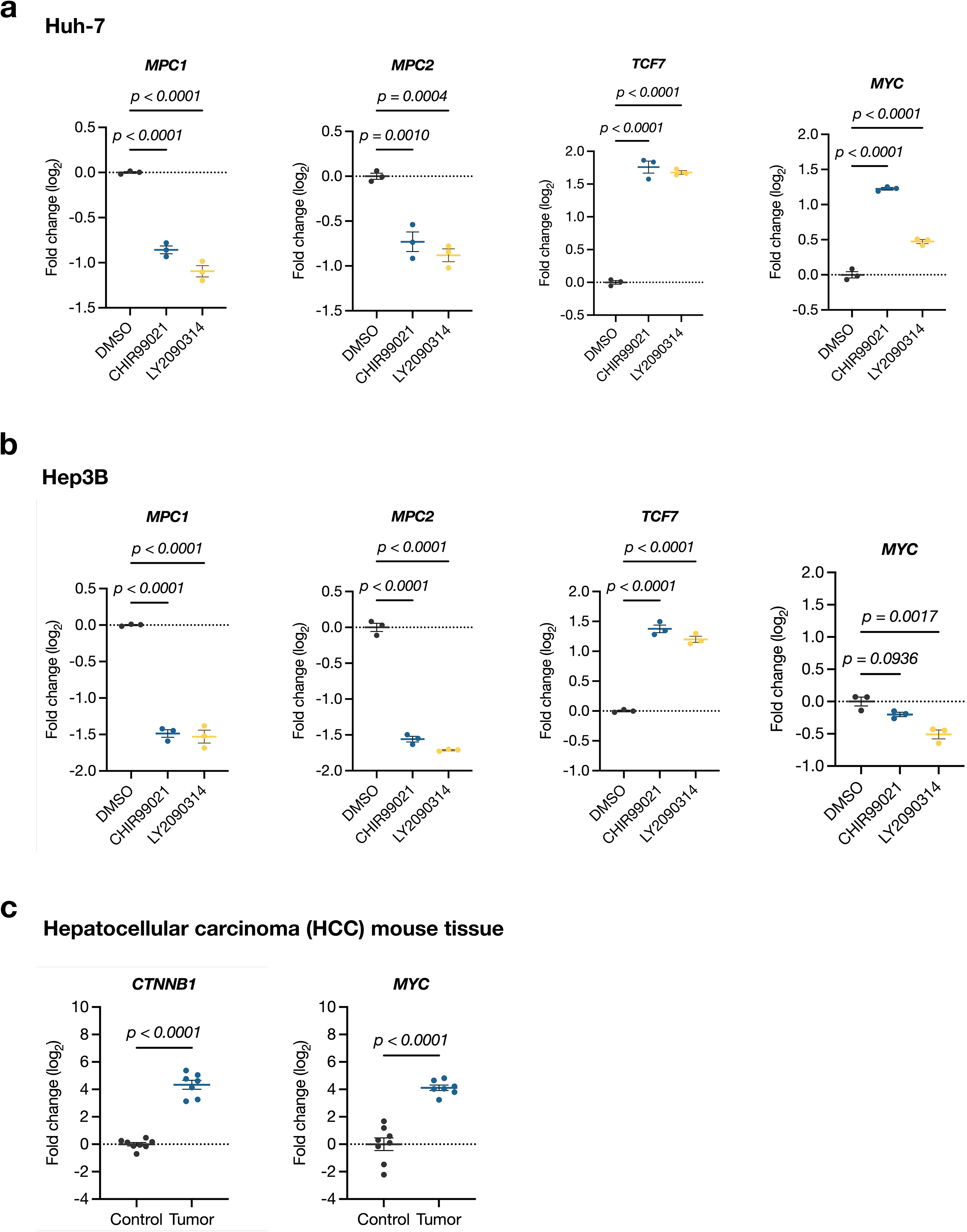
β-catenin activation represses MPC expression in additional liver cancer cell lines, and human *CTNNB1* and *MYC* transgenes are expressed in mouse HCC tumors. **a**,**b**, RT–qPCR of *MPC1*, *MPC2*, *TCF7* and *MYC* in Huh-7 (**a**) and Hep3B (**b**) cells treated with DMSO, 20 µM CHIR99021 or 5 µM LY2090314 for 48 h. Data are log₂ fold change relative to DMSO, mean ± s.e.m. of *n* = 3 independent biological replicates; One-way ANOVA with Dunnett correction. **c**, RT– qPCR of human *CTNNB1* and *MYC* transgene expression in liver tissue from control mice and in tumors from mice injected with Sleeping Beauty transposons encoding human *CTNNB1* and *MYC*. Data are log₂ fold change relative to control, mean ± s.e.m.; *n* = 8 control and 8 tumor samples; two-tailed unpaired Student’s *t*-test. Exact *p* values are shown in each panel.

**Extended Data Fig. 2.**
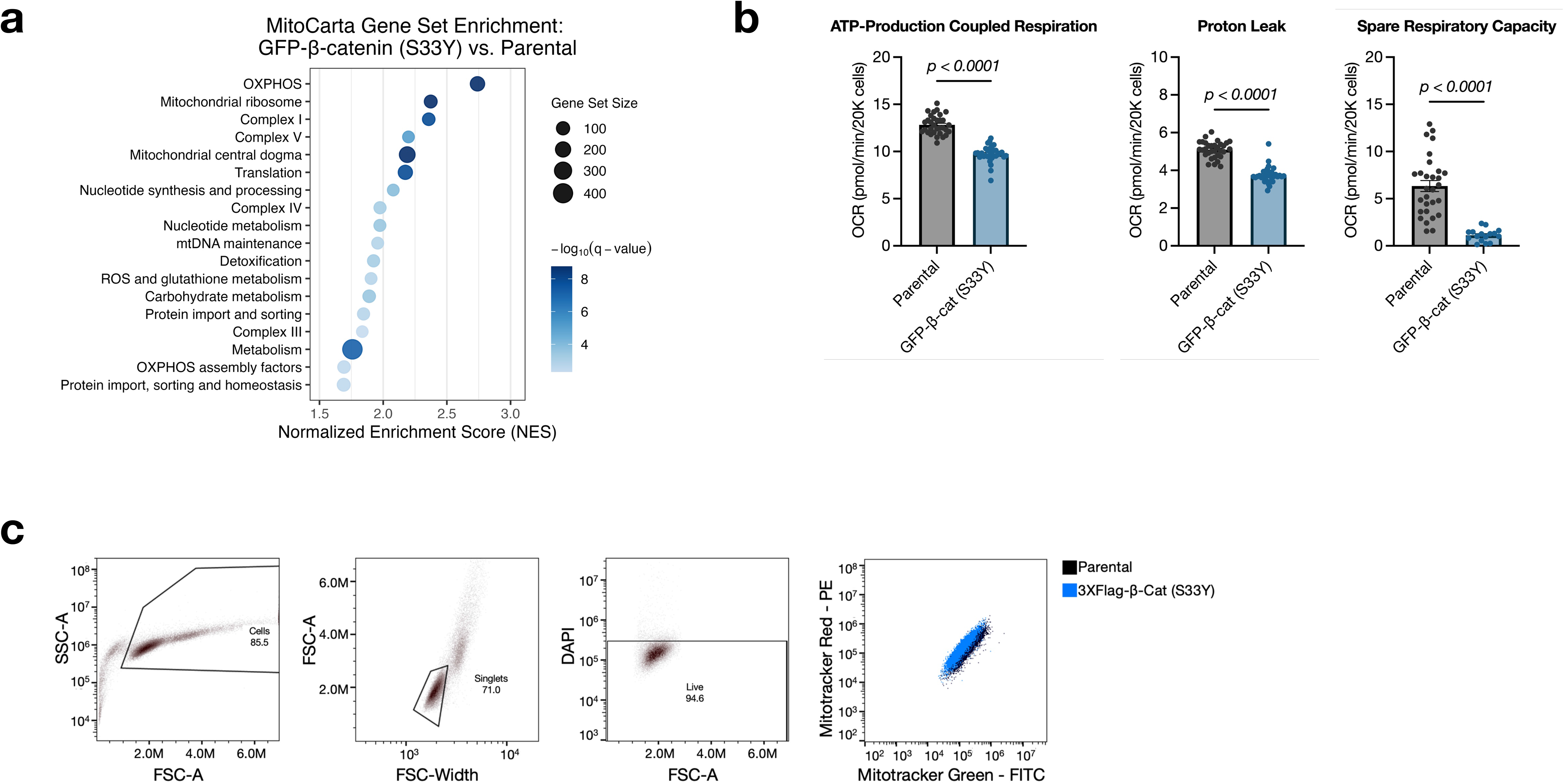
Mitochondrial gene set enrichment, respiratory parameters and gating strategy. **a**, MitoCarta gene set enrichment analysis of RNA sequencing data from β-catenin (S33Y) versus parental JHH7 cells (*n* = 3 independent biological replicates per group), ranked by normalized enrichment score (NES); dot size indicates gene set size and color indicates −log₁₀(*q* value). **b**, ATP-production-coupled respiration, proton leak and spare respiratory capacity derived from the Seahorse experiment shown in Fig. 2c. Data are mean ± s.e.m. of *n* = 4 independent experiments, each comprising 8 technical replicate wells; statistical significance was assessed on the experimental means by two-tailed unpaired Student’s *t*-test. **c**, Gating strategy for the mitochondrial membrane potential measurements in Fig. 2i. Cells were gated on FSC-A/SSC-A, singlets on FSC-A/FSC-width, and live cells by DAPI exclusion. Right, representative MitoTracker Red versus MitoTracker Green plot for parental and β-catenin (S33Y) cells.

**Extended Data Fig. 3.**
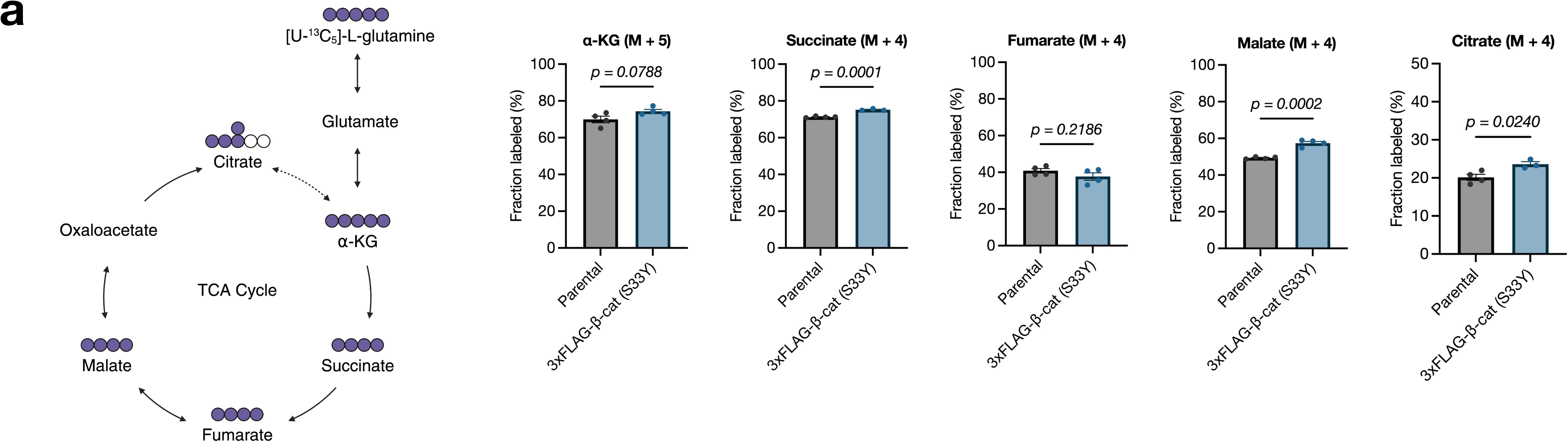
Glutamine labelling of the TCA cycle is largely preserved upon β-catenin activation. **a**, Left, schematic of [U-¹³C₅]-L-glutamine metabolism through glutamate and the TCA cycle. Right, fractional enrichment of M+5 α-ketoglutarate and M+4 succinate, fumarate, malate and citrate following 2 h incubation with [U-¹³C₅]-L-glutamine. Data are mean ± s.e.m. of *n* = 4 independent biological replicates; statistical significance was assessed by two-tailed unpaired Student’s *t*-test. Exact *p* values are shown in each panel.

**Supplementary Table 1a.**
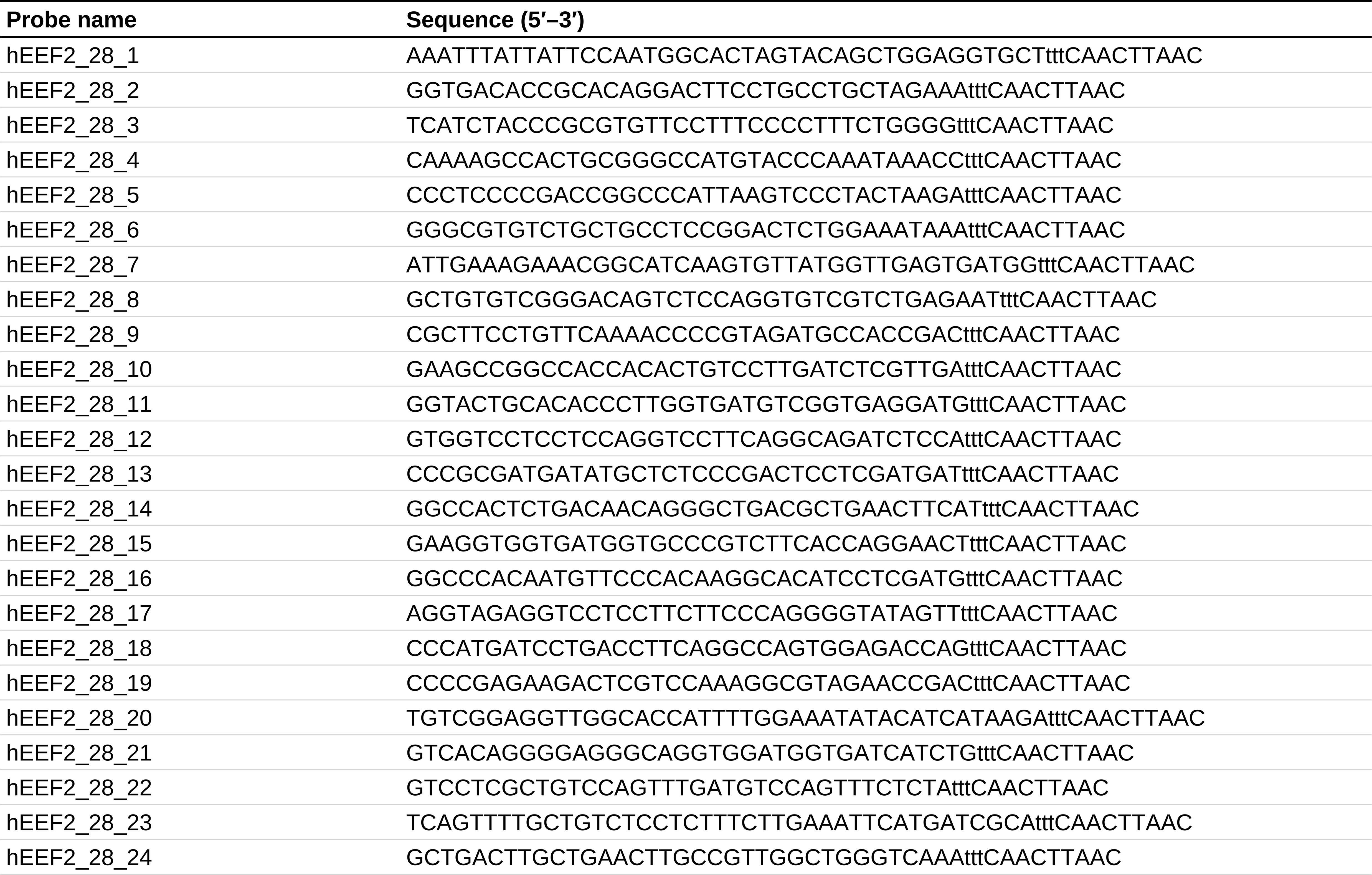
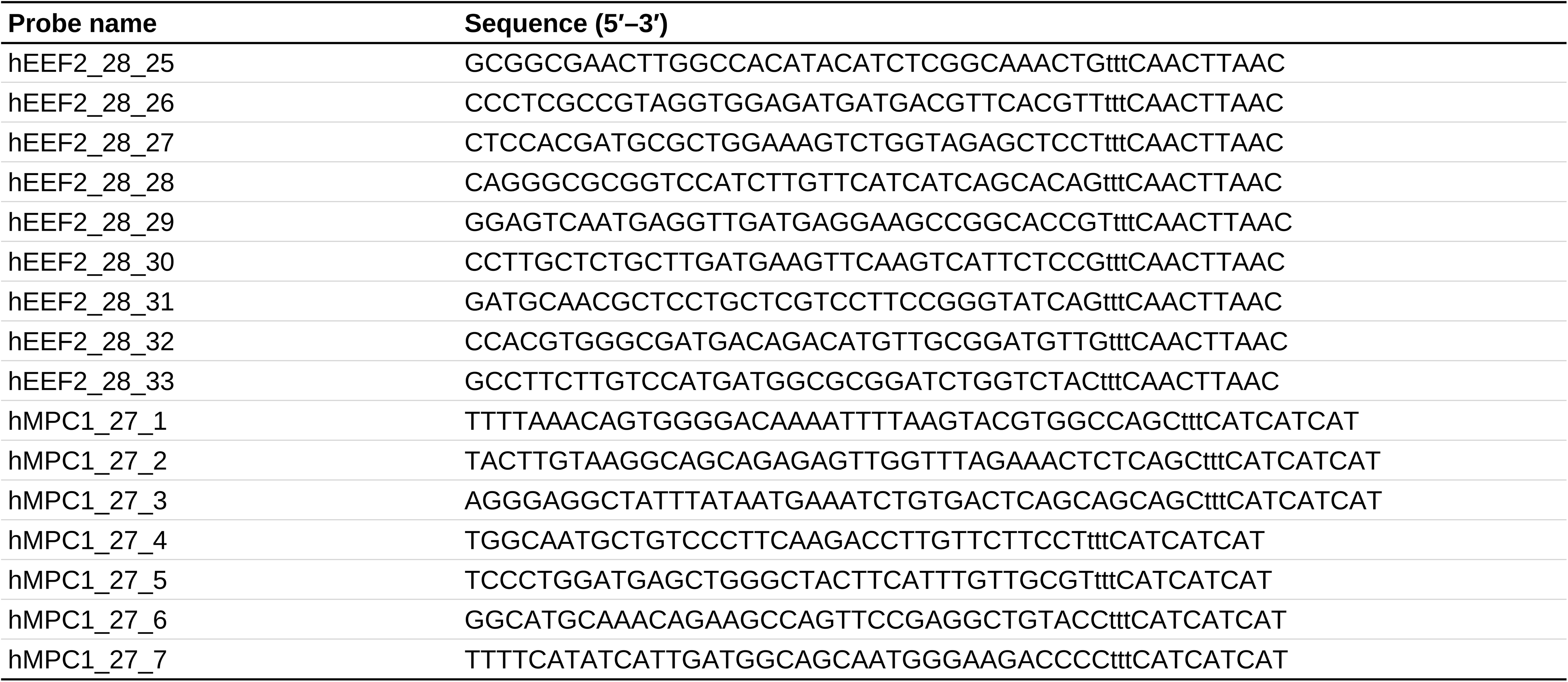
TRoUT-FISH primary probe sequences targeting human MPC1 and EEF2. Oligonucleotide probes used for RNA FISH against MPC1 (7 probes) and the reference gene EEF2 (33 probes). Each probe consists of a target-binding sequence followed by a 3′ extension shared by all probes for that gene (tttCAACTTAAC for EEF2; tttCATCATCAT for MPC1). Sequences are written 5′ to 3′.

**Supplementary Table 1b.**
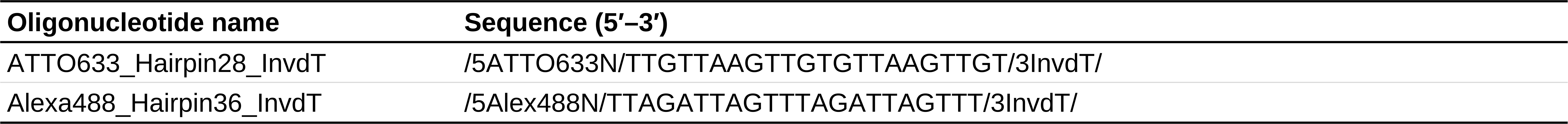
Fluorescently labeled oligonucleotides used for TRoUT-FISH detection. Fluorophore-conjugated oligonucleotides with a 3′ inverted dT (InvdT). Sequences are written 5′ to 3′; modifications are given in slashes.

**Supplementary Table 1c.**
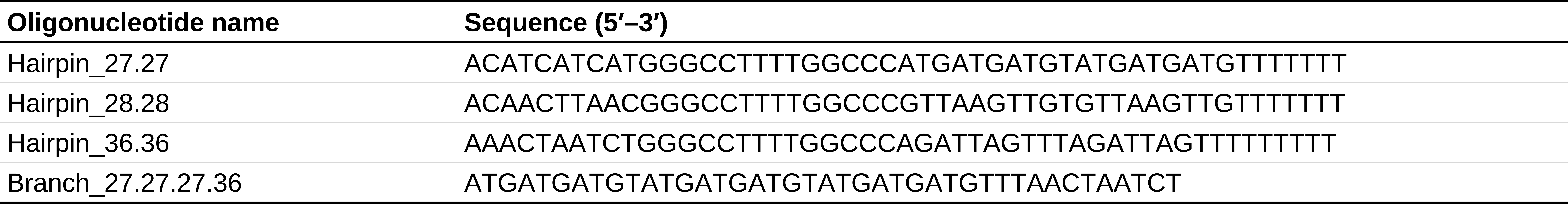
Primer exchange reaction (PER) hairpin and branch oligonucleotide sequences. Hairpin and branch oligonucleotides used for PER-based signal amplification in TRoUT-FISH. Sequences are written 5′ to 3′.

**Supplementary Table 2.**
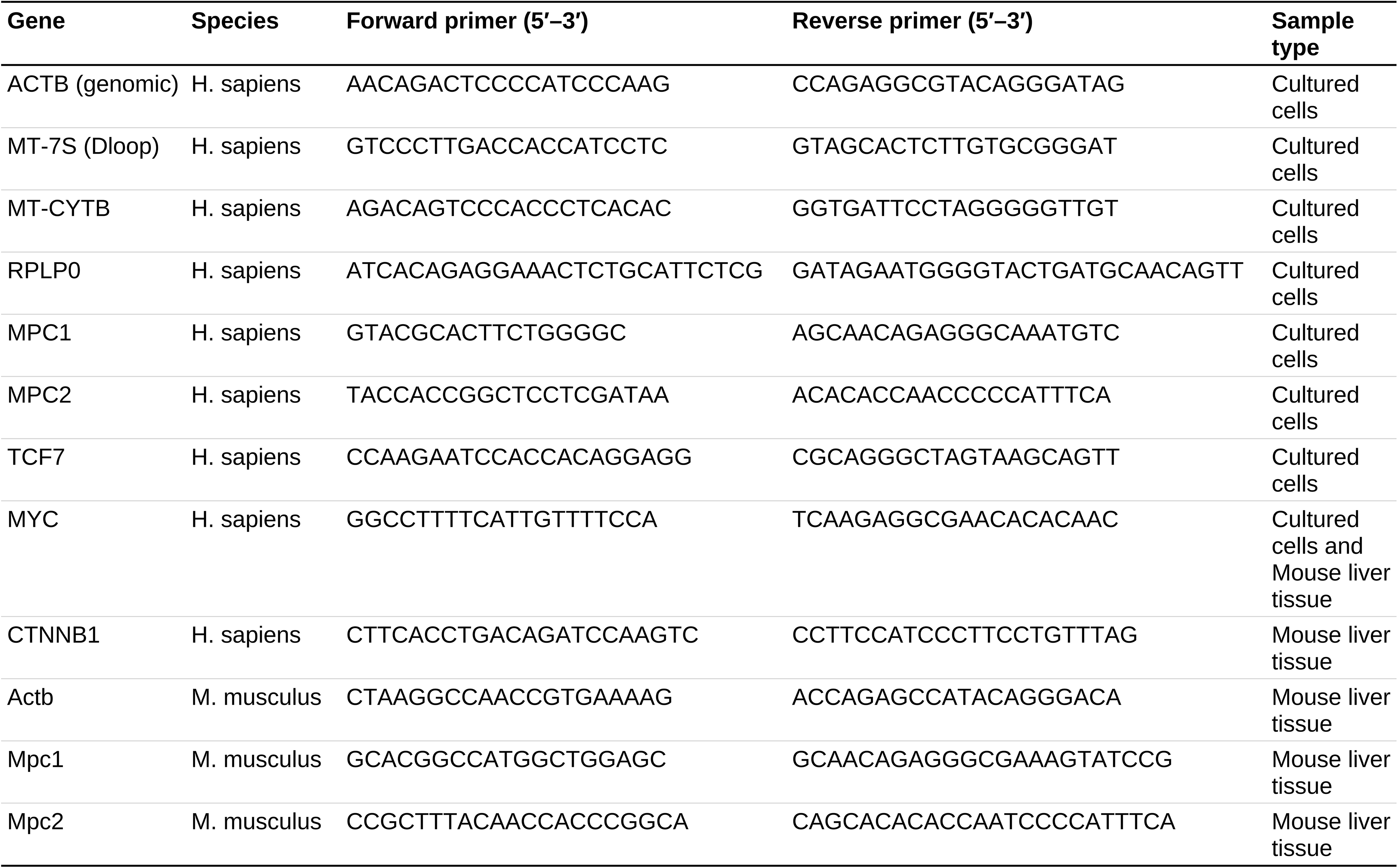
Primer sequences used for RT–qPCR. Primers used for RT–qPCR in cultured cells and in mouse liver tissue. Sequences are written 5′ to 3′.

